# Scalable Hypothalamic Arcuate Neuron Differentiation from Human Pluripotent Stem Cells Suitable for Modeling Metabolic and Reproductive Disorders

**DOI:** 10.1101/2024.06.27.601062

**Authors:** Vukasin M. Jovanovic, Narisu Narisu, Lori L. Bonnycastle, Ravi Tharakan, Kendall T. Mesch, Hannah J. Glover, Tingfen Yan, Neelam Sinha, Chaitali Sen, David Castellano, Shu Yang, Dvir Blivis, Seungmi Ryu, Daniel F. Bennett, Giovanni Rosales-Soto, Jason Inman, Pinar Ormanoglu, Christopher A. LeClair, Menghang Xia, Martin Schneider, Erick O. Hernandez-Ochoa, Michael R. Erdos, Anton Simeonov, Shuibing Chen, Francis S. Collins, Claudia A. Doege, Carlos A. Tristan

## Abstract

The hypothalamus, composed of several nuclei, is essential for maintaining our body’s homeostasis. The arcuate nucleus (ARC), located in the mediobasal hypothalamus, contains neuronal populations with eminent roles in energy and glucose homeostasis as well as reproduction. These neuronal populations are of great interest for translational research. To fulfill this promise, we used a robotic cell culture platform to provide a scalable and chemically defined approach for differentiating human pluripotent stem cells (hPSCs) into pro-opiomelanocortin (POMC), somatostatin (SST), tyrosine hydroxylase (TH) and gonadotropin-releasing hormone (GnRH) neuronal subpopulations with an ARC-like signature. This robust approach is reproducible across several distinct hPSC lines and exhibits a stepwise induction of key ventral diencephalon and ARC markers in transcriptomic profiling experiments. This is further corroborated by direct comparison to human fetal hypothalamus, and the enriched expression of genes implicated in obesity and type 2 diabetes (T2D). Genome-wide chromatin accessibility profiling by ATAC-seq identified accessible regulatory regions that can be utilized to predict candidate enhancers related to metabolic disorders and hypothalamic development. In depth molecular, cellular, and functional experiments unveiled the responsiveness of the hPSC-derived hypothalamic neurons to hormonal stimuli, such as insulin, neuropeptides including kisspeptin, and incretin mimetic drugs such as Exendin-4, highlighting their potential utility as physiologically relevant cellular models for disease studies. In addition, differential glucose and insulin treatments uncovered adaptability within the generated ARC neurons in the dynamic regulation of POMC and insulin receptors. In summary, the establishment of this model represents a novel, chemically defined, and scalable platform for manufacturing large numbers of hypothalamic arcuate neurons and serves as a valuable resource for modeling metabolic and reproductive disorders.

## Introduction

In the mid-1930s, Ernst and Berta Scharrer first proposed that the hypothalamus contains neurosecretory cells ^1^. Numerous physiological functions are regulated by the hypothalamus, such as the maintenance of body temperature, the regulation of hormone secretion and the circadian rhythm-based sleep-wake cycle ^2^. The hypothalamic arcuate nucleus (ARC) is involved in the regulation of energy balance, reproduction, and neuroendocrine control of growth hormone and prolactin release ^3,4^. Thus, impaired function of ARC neurons contributes to metabolic disorders such as obesity and type 2 diabetes (T2D) ^5–7^.

During early brain development, the ARC emerges from the ventral diencephalon and is marked by the expression of specific transcription factors, such as the homeodomain transcription factors RAX and NKX2.1 ^8^. Neurogenesis unfolds through the transcriptional activation of ASCL1, NEUROD1, NEUROG3, DLX1/2/5, ARX, and OTP as these progenitors differentiate and transition into neuroblasts ^8^. Subsequently, distinct neuronal populations form within the ARC. Among them are the anorectic POMC neurons, essential for regulating feeding behavior and energy balance ^9^.

POMC neurons respond to postprandially secreted hormones such as insulin, elevated levels of circulating glucose and the adipose secreted hormone leptin, all of which result in appetite suppression. In contrast, orexigenic Agouti-related peptide (AgRP) neurons within the ARC, specified later in hypothalamic development ^10^, counteract activation of POMC neurons to promote appetite. Additional ARC neuronal populations include GnRH neurons ^11^, kisspeptin (KISS1) neurons that stimulate GnRH neurons, and tuberoinfundibular dopaminergic neurons that regulate prolactin release ^12,13^ as well as several neuronal subtypes, with unique combinations of neuropeptides and transcription factors, whose functions have yet to be described ^10,14^.

Human hypothalamic tissue in living persons is inaccessible for direct functional interrogation. Thus, the ability to efficiently and controllably produce a large number of human hypothalamic ARC neurons *in vitro* would be a valuable resource for investigating the molecular causes of metabolic diseases linked to the hypothalamus, such as obesity and T2D. Moreover, this resource could facilitate the development of therapeutics targeting these and other metabolic disorders. However, previously published protocols for differentiating human pluripotent stem cells (hPSC) to hypothalamic neurons have not addressed scalability ^15–18^.

In this study, we report a scalable, efficient, and controlled approach for the differentiation of hPSCs into populations of hypothalamic ARC neurons. These neurons were comprehensively validated on a cellular and molecular level. They display expression of pertinent ARC markers, as well as functional response to humoral and hormonal stimuli. The scalability of this approach is demonstrated by adapting the protocol to an established robotic cell culture platform ^19,20^ and their translational applicability is documented by the generation of dose-response curves in targeted drug challenges. The data presented in this study will serve as a valuable resource, and the protocol can facilitate at-scale acquisition of previously limited cell types for further studies of ARC-associated disorders.

## Results

### A Highly Efficient Method for Large-Scale Automated Production of Human Hypothalamic Neurons

hPSCs were cultured under feeder-free, chemically defined conditions using E8 medium and vitronectin as a coating substrate. To ensure stress-free cell expansion, the cells underwent EDTA-mediated passaging, followed by treatment with the CEPT small molecule cocktail for 24 h to enhance cell viability ^21^. On day −1, 20,000 hPSCs/cm^2^ were seeded and cultured for 24 h in E8 medium with CEPT. The following day (day 0), the media was changed to Arc-1 medium composed of E6/N2/B27 without vitamin A. Previous reports to direct cells towards a hypothalamic progenitor stage have involved either dual SMAD inhibition combined with activation of Sonic Hedgehog (SHH) signaling, with subsequent inhibition of Notch signaling or dual SMAD and WNT inhibition followed by activation of SHH ^16,22,17^. Here, distinct from previous protocols, early and simultaneous dual SMAD and WNT inhibition combined with SHH activation was accomplished using base medium supplemented with BMP inhibitor LDN193189, TGF-B inhibitor A83-01, WNT inhibitor XAV939 and SHH activator SAG21K in the presence of recombinant proteins (DLL1, JAGG1, LIF and CNTF) designed to simultaneously support neural induction and patterning into ventral diencephalon progenitors. Cells were kept in Arc-1 medium for 7 days to reach a fully confluent monolayer (Fig. 1A).

**Figure 1.**
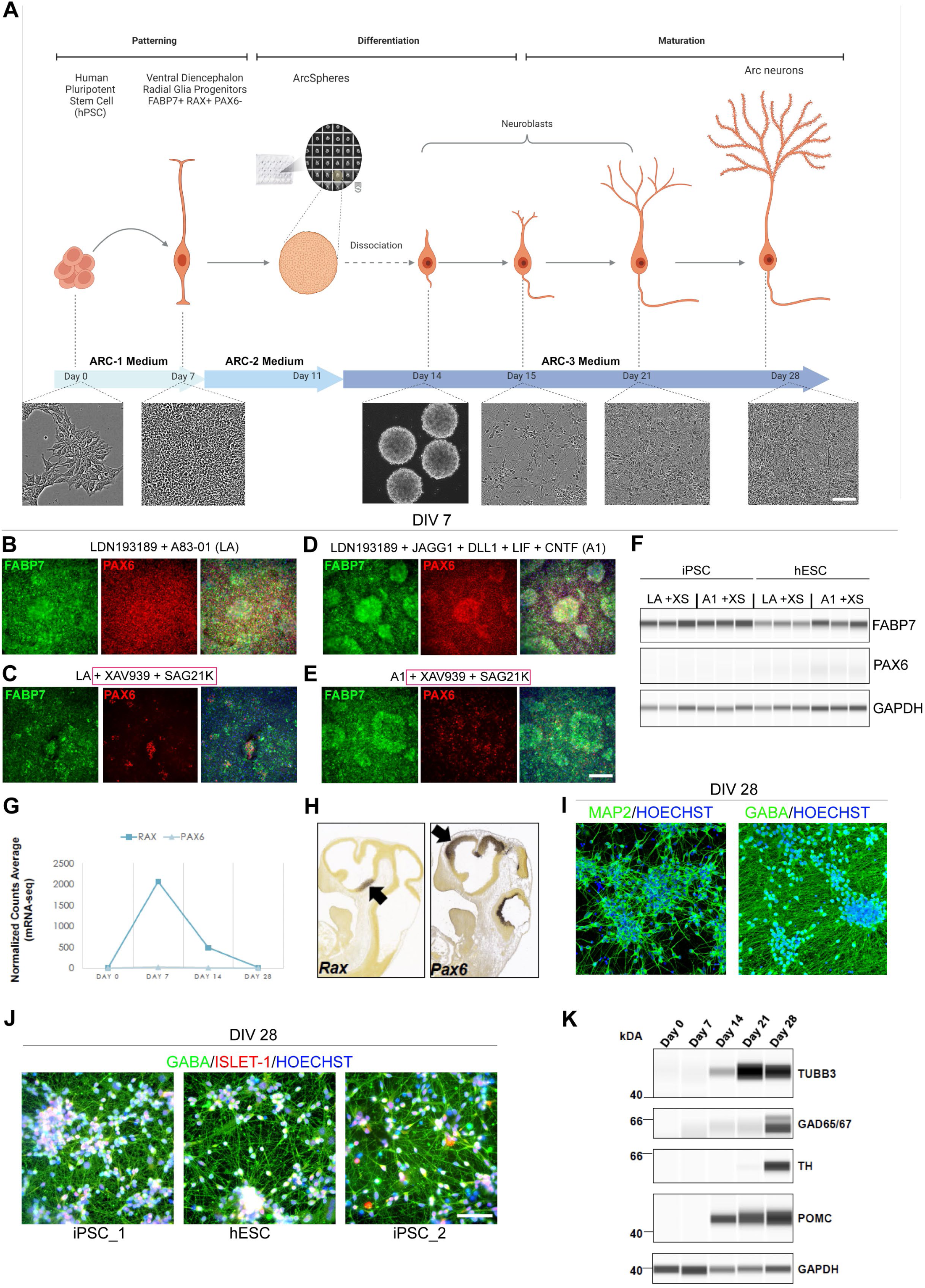
Directed Differentiation of hPSCs into Ventral Diencephalon and Human Hypothalamic Arcuate Neurons. (A) Schematic overview of the differentiation protocol with phase contrast images representing the appearance of the cells at day 0, 7, 15, 21, and 28, and ArcSpheres at day 14. (B-E) Representative images of day 7 cells (WA09) immunolabeled for PAX6 and FABP7 in four different treatment conditions. Note the consistent downregulation of PAX6 with the addition of XAV939 and SAG21K to the differentiation medium. (F) Western blot analysis of FABP7 and PAX6 in both treatment conditions including XAV939 and SAG21K. Note that FABP7 expression was maintained. (G) mRNA-seq shows *RAX* and *PAX6* mRNA transcript expression during differentiation of hESC (WA09). Note that *PAX6* mRNA was not upregulated at any profiled time point during the differentiation. (H) *In situ* hybridization of *Rax* and *Pax6* mRNA during the neurogenesis stage of mouse brain development (E11.5). Note that PAX6 is expressed in the dorsal forebrain and the midbrain and is absent from the ventral forebrain. RAX transcripts are present in the ventral diencephalon, neural tube territory that gives rise to the retina, optic stalk, and ventral hypothalamus. Images were downloaded from the Allen Brain Atlas. (I) Representative images of DIV 28 cultures labeled for MAP2 and GABA. HOECHST was used for DNA/nuclear counterstain. Note that virtually all cells present neuronal identity (MAP2) and inhibitory neuron (GABA) marker expression. (J) Neurons derived from three independent hPSC lines (LiPSC-GR 1.1, WA09, and NCRM-5 respectively) at DIV 28 immunolabeled for ISL-1 and GABA. Note the consistency of differentiation across the independent cell lines. (K) Time-course Western blot analysis of differentiating cells shows a gradient of upregulation of neuronal marker TUBB3 (TUJ1), GAD65/67 inhibitory neuron markers, and POMC arcuate neuron marker starting at DIV 14. The dopaminergic arcuate neuron population is developed later, with TH protein expression first detected on DIV21 and upregulated strongly at DIV 28. Scale bars 100 µm.

We previously reported that hPSCs exposed to recombinant protein modulators of JAK/STAT and NOTCH signaling in addition to BMP inhibition (LDN193189 + JAGG1 + DLL1 + LIF + CNTF) directly differentiated into radial glial cells expressing FABP7 and PAX6, a marker of dorsal neural tube progenitors ^23,20^. To identify culture conditions that promote the generation of ventral diencephalon progenitors, FABP7 and PAX6 expression was assessed on day 7 after treatment in one of four distinct conditions: LDN 193189 + A83-01 (LA) (Fig. 1B), LA + XAV939 + SAG21K (XS) (Fig. 1C), LDN193189 + JAGG1 + DLL1 + LIF + CNTF (A1) (Fig. 1D), or A1 + XAV939 + SAG21K (Fig. 1E). In agreement with directing the cells towards the ventral fate, we observed a decrease of PAX6 expression in both treatment groups containing XAV939 and SAG21K small molecules, while FABP7 expression was maintained indicating that cells were directed towards a neural progenitor stage and ventralized under these culture conditions. To ascertain robustness of this culture condition, these findings were further validated by Western blot in two independent hPSC lines (WA09 human embryonic stem cells (hESC) and LiPSC-GR 1.1 hiPSC (Fig. 1F). A1 treatment induced strong FABP7 expression in both cell lines (Fig. 1F), as previously described ^20^. Importantly, *PAX6* mRNA expression after Arc-1 medium treatment was also not observed in an orthogonal mRNA-seq time-course experiment, while *RAX*, an essential regulator of hypothalamus specification ^24^, was strongly induced at day 7 (Fig. 1G and 1H). These ventral diencephalon progenitors were then placed into AggreWell plates to form neurospheres (ArcSpheres) and maintained in Arc-2 medium as free-floating spheres to maximize neurogenesis until day 11 (Fig. 1A). Since WNT inhibition and Activin/Nodal signaling have been associated with hypothalamic development ^25,26^, Arc-2 medium was composed of DMEM/F12 with N2 and B27 without vitamin A, supplemented with XAV939, Activin A, BDNF, GDNF, Ascorbic Acid and db-CAMP. On day 11, Arc-2 medium was replaced with Arc-3 medium; a standard neuronal maturation medium comprised of DMEM/F12 with N2 and complete B27, supplemented with BDNF, GDNF, Ascorbic Acid, db-CAMP, with addition of gamma-secretase inhibitor dibenzazepine (DBZ) ^27^. Spheres were dissociated on day 14 and cultured as a monolayer in Arc-3 medium for an additional two weeks. On day 28, virtually all cells expressed the neuronal marker MAP2, and the majority of neurons expressed inhibitory marker gamma-aminobutyric acid (GABA) (Fig. 1I). LIM-homeodomain transcription factor Islet 1 (ISL1) is expressed in the developing hypothalamus immediately before the onset of POMC expression and from then on ISL1 and POMC are coexpressed ^28^. Neurons derived from three independent hPSC lines exhibited ISL1 expression at day 28 (Fig. 1J). Recently, single-cell RNA sequencing experiments have revealed heterogeneity among POMC expressing neurons ^29,30^. Furthermore, in the hypothalamus reference dataset (HypoMap), multi-level clustering of cell populations generated seven levels of clusters, within these, one cluster level (C7) included cell populations expressing POMC and GAD65/67, while five cluster levels (C25, C66, C185, C286 and C465) contained cell populations expressing POMC, GAD65/67 and TH ^31^. To further validate neuronal and hypothalamic arcuate identity of the derived cells, we performed time-course Western blot experiments and detected the induction of POMC at day 14 that correlated to the onset of expression of beta III tubulin (TUBB3) (Fig. 1K). By day 28, these hPSC-derived neurons strongly expressed tyrosine-hydroxylase (TH), as well as glutamate decarboxylase enzymes (GAD65/67) which play roles in GABA synthesis (Fig. 1K).

Next, we performed high-content imaging of the neuronal cultures derived from three independent hPSC lines (Fig. 2A) by generating entire well images of POMC, TH, somatostatin (SST), and GAD65/67 expression markers (Fig 2B). Unbiased high-content imaging analysis allowed for quantification of Hoechst^+^ nuclei and specific markers of interest (Fig. 2C). This analysis demonstrated a degree of heterogeneity and consistency in the type and number of neurons expressing specific markers across three independent hPSC lines, with POMC^+^ ranging from 44-56% (Fig. 2D), dopaminergic TH^+^ neurons ranging from 12-15% (Fig. 2E), and SST^+^ neurons ranging from 3-6% (Fig. 2F) when normalized to Hoechst^+^ nuclei.

**Figure 2.**
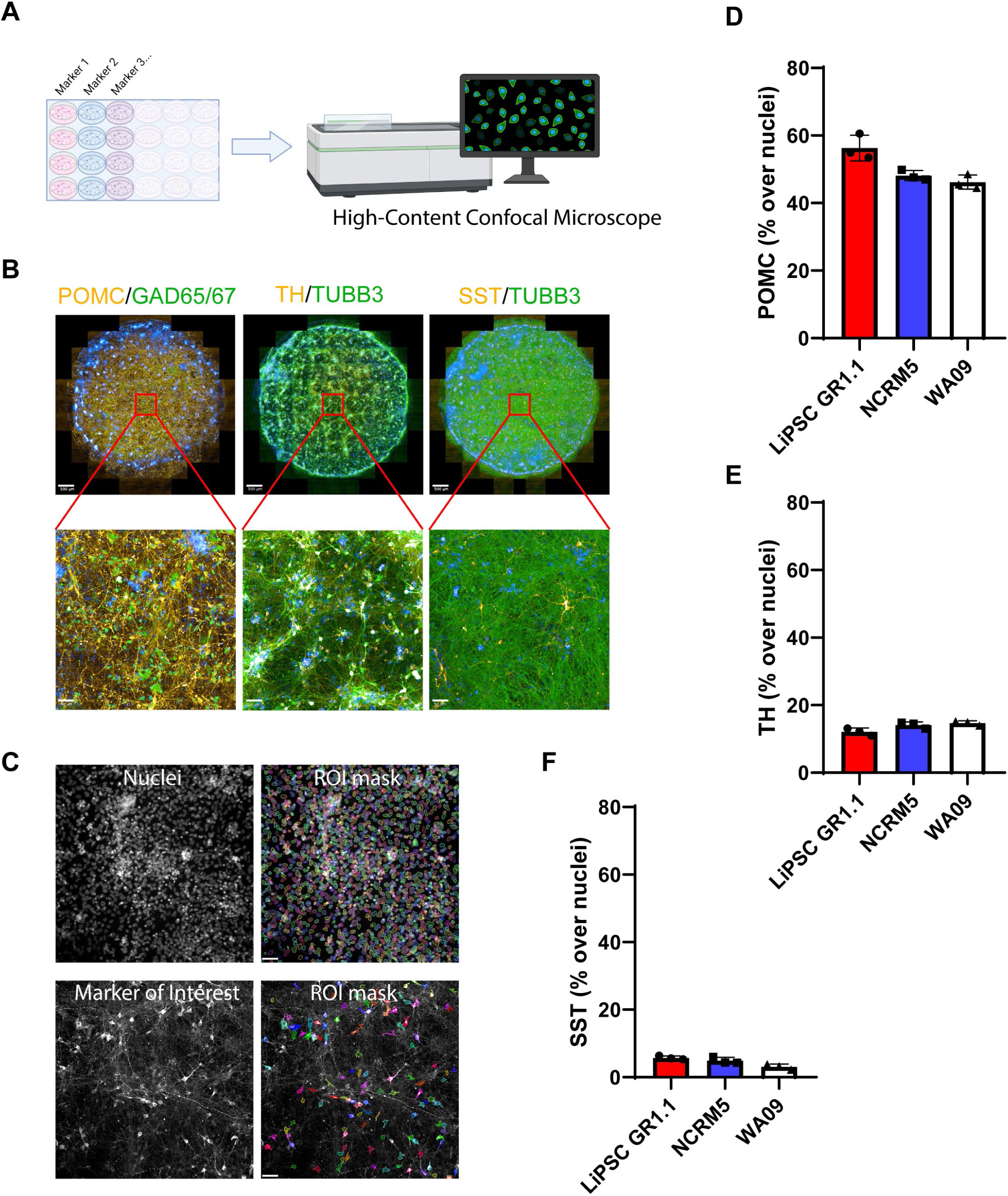
High-content Imaging Analysis of Hypothalamic Neuron Cultures. (A) Schematic overview of the high-content imaging design and the OperaPhenix imager. (B) Whole-well images (24-well plate/0.5cm^2^ area per well/89 images) of DIV 28 neurons capturing the consistent expression of POMC, GAD65/67, TH, TUBB3, and SST across the cultures. (C) Region of interest (ROI) masks capturing nuclei and the neuronal marker of interest (TH marker segregation represented as an example). Unbiased batch image analysis was performed in Columbus software (Revvity), using a set of predefined parameters. (D–F) Quantification of POMC, TH, and SST percentage per well. For each marker, three replicate wells were quantified, and each data point represents one well. Scale bars 500 µm (B – upper row), 50 µm (B -bottom row and C). Hypothalamic neurons derived from LiPSC-GR1.1 were used for high-content imaging experiments.

Given the reproducibility and robustness of this differentiation protocol across multiple hPSC lines, we reasoned that fully automating the entire process on a non-stop robotic cell culture platform for scale-up and biomanufacturing could provide an unlimited source of hypothalamic arcuate neurons (Fig. S1A). To this end, this protocol was automated using the CompacT SelecT platform where three million hPSCs were differentiated in three T-175 flasks to generate up to 105 million hypothalamic neurons by day 14. At day 14 hypothalamic neurons can be cryopreserved or undergo a 2-week maturation process in the Arc-3 medium before assaying (Fig. S1B). Upon thawing, cryopreserved hypothalamic neurons matured in Arc-3 medium until day 28 display ARC markers and are suitable for studies of ARC-associated disorders.

### Time-course mRNA-seq Analysis Confirms Hypothalamic Identity of Differentiated Neurons

*In vivo* development of the human hypothalamus is guided by complex transcription factor programs that shape the molecular architecture of the region into separable nuclei populated by distinct neuronal populations ^8^. NKX2.1 and RAX positive hypothalamic progenitors undergo basic-helix loop helix transcription factor - ASCL1 mediated neurogenesis to transition into postmitotic arcuate neuron precursor subpopulations. Further specification of these postmitotic precursors into arcuate neuron subpopulations is driven by combinatorial induction of TBX3, OTP and DLX1/2/5 ^32^.

To capture the stepwise transcriptome changes throughout the differentiation of hPSCs into hypothalamic neurons, we profiled a total of 44 RNA samples isolated from three independent hPSC lines (LiPSC-GR 1.1, NCRM5, and WA09) at days 0, 7, 14, 21, and 28. Principal component analysis (PCA) showed clustering of samples according to the time points profiled during differentiation (Fig. 3A), demonstrating the reproducibility of our protocol and pipeline. We used the top upregulated genes at each time point to map the stepwise acquisition of hypothalamus identity throughout the *in vitro* differentiation process for all three hPSC lines tested. The hypothalamus specification of hPSCs was reflected by early induction of *RAX* and *NKX2.1* in day 7 progenitors, followed by an upregulation of *ASCL1, DLX1/2/5* on day 14, and finally by day 28 strong expression of *TH*, *SST* and *POMC* (Fig. 3B). To characterize the presence of major neuronal subpopulations, we analyzed the time-course expression of several markers, *SLC32A1* for inhibitory GABA neurons, *TH* for dopamine neurons, *HDC* for histamine-producing neurons, *SLC17A5* for excitatory neurons, and *SLC6A4* for serotonin neurons, as described in Herb et al. ^14^ (Fig. 3C). This analysis demonstrated that transcripts of *SLC32A1* and *TH* were most abundantly upregulated and detected by day 28, with some evidence of *HDC* detection, albeit at lower levels. Low upregulation of *SLC17A5* was detected only in hypothalamic neurons derived from one out of three cell lines (WA09 hESC) (Fig. 3C). Collectively, these findings suggest a predominant representation of inhibitory GABA neurons in our cultures. This is consistent with our immunocytochemistry findings (Fig. 1J), which also demonstrated that some GABA neurons co-express SST (Fig. S2A) and Calbindin (Fig. S2B) ^14^.

**Figure 3.**
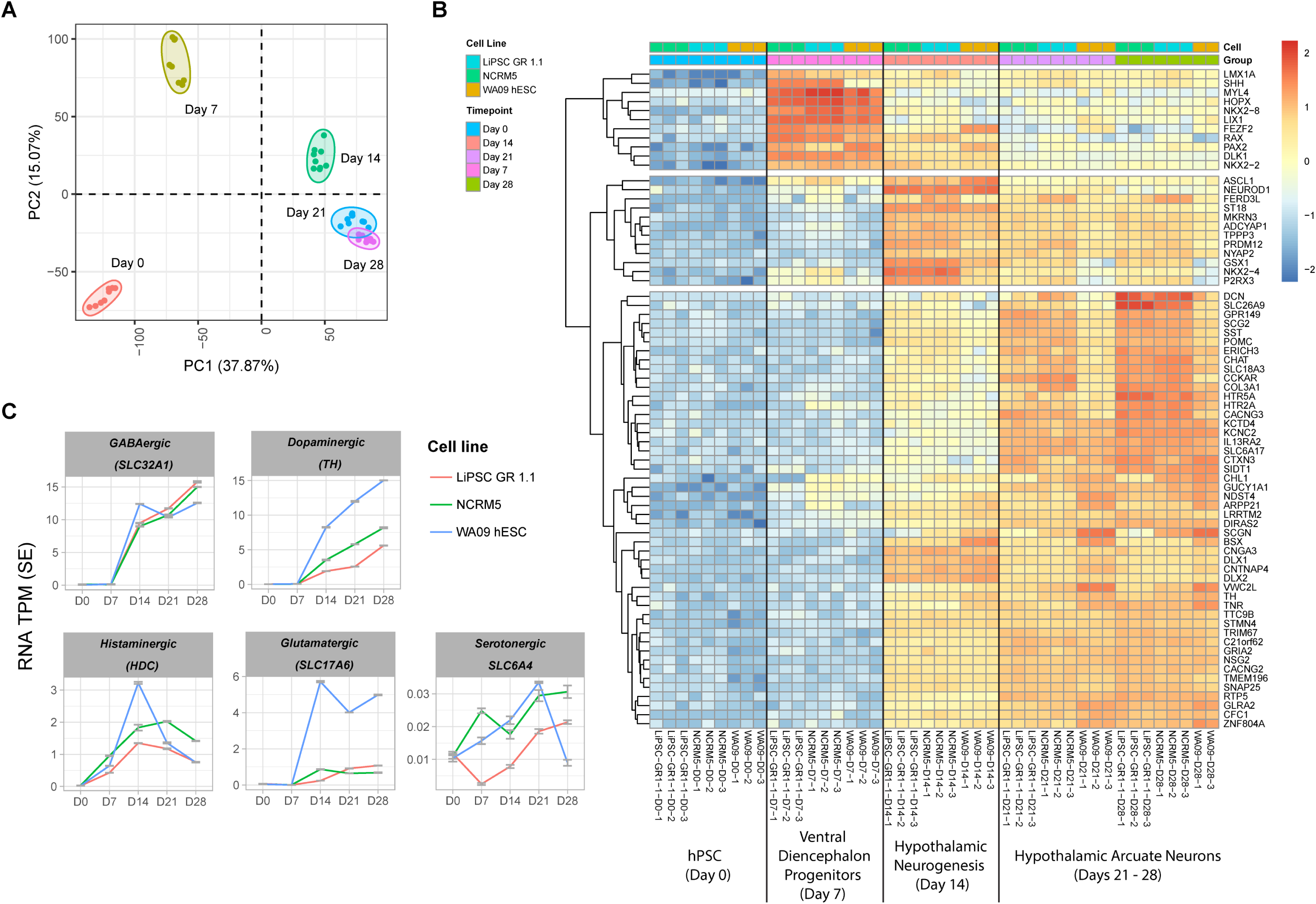
Time-course Transcriptomic Analysis of hPSC Differentiation into Ventral Diencephalon and Hypothalamic Arcuate Neurons. (A) PCA of a time-course mRNA-seq experiment including samples from three independent cell lines. Note the clustering of the samples according to the time-point of differentiation, suggesting the same developmental trajectory between samples from different cell lines. (B) The heatmap represents the top 20 gene markers upregulated (log2FC) at days 7, 14, and 21, and the top 60 gene markers upregulated at day 28, relative to DIV 0 (hPSC). Due to the overlap of top markers between the time points, the total number of genes represented on the heatmap is 71. Note the stepwise upregulation of ventral diencephalon markers RAX, NKX2.1, ASCL1, DLX1, and ARX, as well as arcuate nucleus markers ISL1, POMC, SST, and TH. N = 3 independent hPSC lines differentiation experiments for all datasets presented (LiPSC-GR 1.1, WA09, NCRM5).

Next, we sought to further validate hypothalamic identity of our neurons by performing gene pathway enrichment analyses and comparing it to publicly available datasets from previous studies. Indeed, when we used the top 100 genes upregulated at day 28 as compared to day 0 (log_2_fc), we found strong association to human hypothalamus samples in the GTEx and Human Gene Atlas databases (Fig. S3A, S3B). Furthermore, circadian entrainment was the top functional ortholog feature in the KEGG database query (Fig. S3C).

To better understand how our cells differ from those derived using a previously published iPSC-differentiation method we employed Remove Unwanted Variation sequencing (RUV-seq) normalization and performed a PCA analysis that demonstrated our iPSC-derived hypothalamic neurons clustered separately from those derived in a previous study (Fig. S3D) ^15^. To investigate the distinct transcriptomic signatures among various arcuate nucleus neuronal subpopulations, we drew upon a recent study that profiled over one hundred thousand single cells throughout the development of the human hypothalamus and constructed a heatmap featuring the top 10 markers expressed across 6 different neuronal subpopulations in the human arcuate nucleus ^10^. We found significant enrichment of the markers related to *POMC* and *SST* neuron subpopulations (middle cluster) in our dataset (Fig. S3E).

As demonstrated by high-content image analysis and the time-course transcriptome profiling above, the majority of neurons at the final timepoint of differentiation are of ARC identity (Fig. S4A and S4C). However, we observed a low level of *AgRP* mRNA (Fig. S4C), a known marker of the appetite promoting ARC neuron subpopulation ^33^. Notably, this was not due to delayed progenitor differentiation, as markers of proliferative neural progenitors such as *NES, SOX2, FABP7 and MKI67* were downregulated by day 21 and 28 (Fig. S4B). This is further supported by the detection of transcripts previously described as marker genes for the dorso-medial (*GPR50*) and ventro-medial (*NPTX2*) hypothalamus ^14^ (Fig. S4D) which are two nuclei in close proximity to the arcuate nucleus (Fig. S4A). There was little to no evidence of marker genes representing other hypothalamic nuclei (Fig. S4E – S4F). In summary, our transcriptomic analysis and validation methods confirm that the differentiation protocol robustly and reproducibly generates hypothalamic neurons, predominantly of ARC identity, with specific enrichment of inhibitory GABA neurons and minimal evidence of other hypothalamic nuclei marker genes.

### Time Course Analysis and Enrichment of Disease-Relevant Genes in hPSC-Derived Hypothalamic Neurons

To assess the differentiation stage of our hPSC-derived hypothalamic neurons, we first performed pairwise correlation analyses across different stages of hPSC differentiation and revealed dynamic transcriptome changes that stabilized by day 21 (Fig. 4A). We also compared day 0 with each of the other profiled stages (day 7, 14, 21 and 28) to identify up-regulated genes relative to day 0. Using a stringent threshold of FDR<1^e-6^ and fold change > 4, we found over 6000 upregulated genes by days 21 and 28. (Fig. S5A). Top upregulated pathways at day 7 included c TGF-ß and Hedgehog signaling pathways, corresponding to ventral diencephalon patterning, while synaptic vesicle cycle, circadian entrainment, and GnRH secretion were upregulated at later time points (day 14, 21, and 28), suggesting hypothalamus neuronal specification (Fig. S5B). Downregulated pathways at later time points correlated with cells exiting the cell cycle (Fig. S5C). A stepwise comparison between day 7 and day 14 provided intriguing insights, outlining the onset of hypothalamic neurogenesis in the upregulated pathways and the downregulation of neural progenitor, and patterning related pathways (TGF-ß, Hedgehog, MAPK, Hippo, Rap1, and Ras signaling) (Fig. S5 D-E).

**Figure 4.**
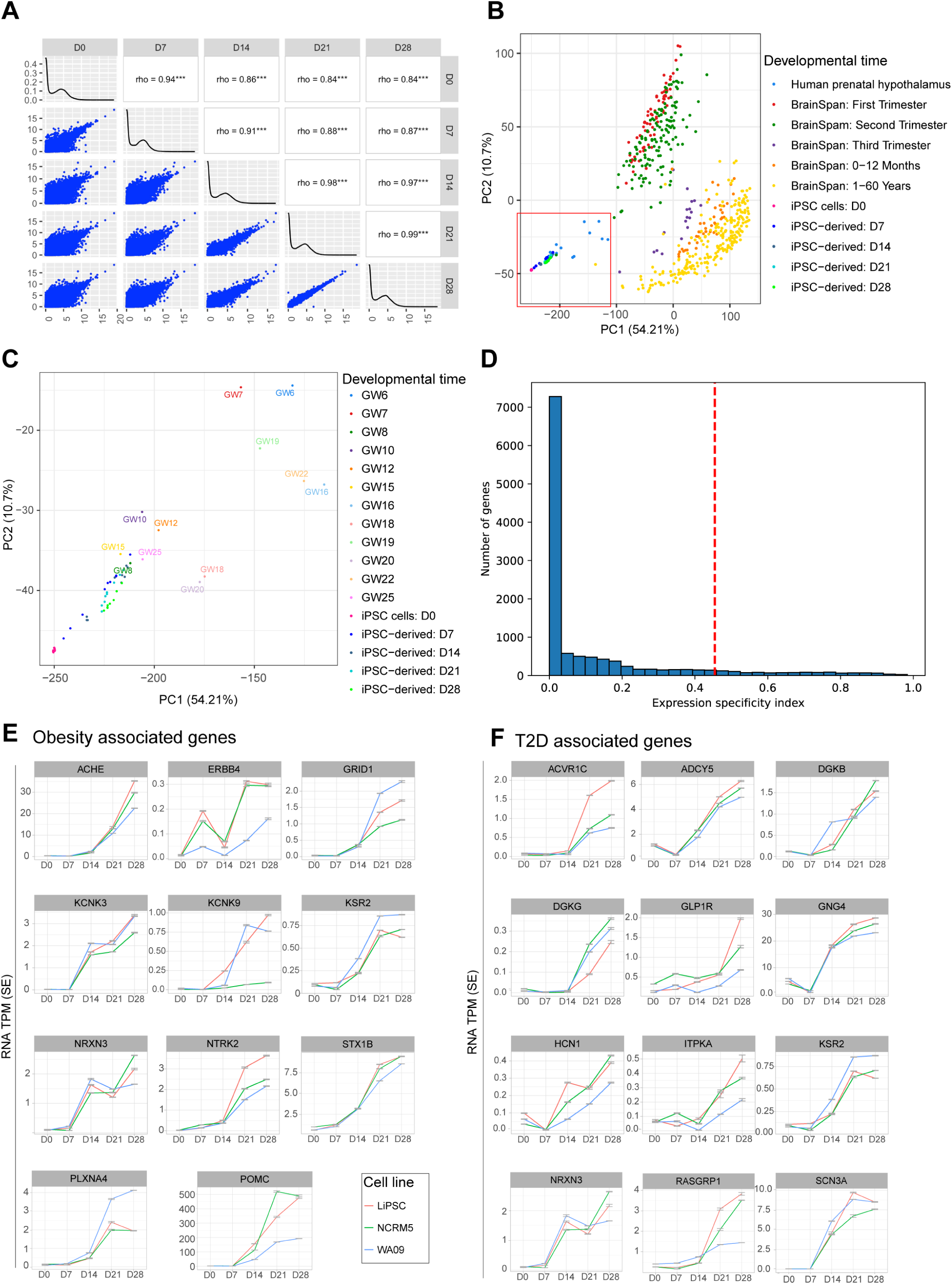
Enrichment of Obesity and T2D Associated Genes in hPSC-Derived Hypothalamic Neurons. (A) Pairwise correlations of gene expression across hPSC differentiation stages. (B-C) Comparison of iPSC-derived neuronal cells with tissues from different brain regions sourced from Brainspan, Spatio-temporal transcriptome of the human brain, and the datasets from Herb et al., Siletti et al, and Zhou et al. studies. Note that (B) specifically includes hPSC-derived and primary hypothalamus samples. (D) Distribution of gene expression specificity index (ESI) at DIV 28. ESI values were calculated using normalized expression data of cells across various developmental stages. Genes located to the right of the vertical red line represent the top 10 percentile of genes with the highest ESI at DIV 28. (E-F) Illustrative examples of genes specific to DIV 28 compared to other developmental stages. This list encompasses monogenic and GWAS obesity genes, as well as T2D associated genes meeting the same criteria. These genes contribute significantly to the enrichment of relevant pathways outlined in Figure S5. Further details can be found in the supplementary figures. N = 3 independent hPSC lines differentiation experiments for all datasets presented (LiPSC-GR 1.1, WA09, NCRM5).

Next, we compared transcriptomes across hPSC differentiation to hypothalamic neurons with 1) those obtained from various human brain regions across different developmental stages (BrainSpan atlas ^34^) and 2) fetal samples from different stages of human hypothalamus development ^10,14,35,36^.

The BrainSpan atlas integrated spatio-temporal dataset of samples from the cortex, amygdala, cerebellum, hippocampus, mediobasal thalamus, and striatum, but not the hypothalamus (Fig. 4B). These samples were collected during the first trimester (red dots), second trimester (green dots), and third trimester (purple dots) of human brain development, as well as during the postnatal period (0-12 months, orange dots) and adulthood (1-60 years, yellow dots) (Fig. 4A). Brainspan samples contain mixed populations of neuronal and non-neuronal tissue. Prenatal hypothalamus samples were from gestational weeks (GW) 6 through 25, marking the first and second trimester of hypothalamus development (Fig. 4B; light blue dots within the red box). PCA analysis illustrated the time-course progression of our iPSC-derived samples towards human fetal prenatal hypothalamus (Fig. 4B and 4C).

Genome-wide association studies (GWAS) have identified hundreds of genomic loci associated with risk for T2D and obesity. Thus, we asked whether any of the associated genes are expressed in our hPSC-derived hypothalamic neurons. To this end, we determined the expression specificity index (ESI), which measures the likelihood that a gene is specifically expressed at the given differentiation time point ^35^. We identified the top 1216 genes with the highest ESI at Day 28 (Fig. 4D; on the right of dashed line and Supplementary Table 2) and compared this set of genes with a group of previously described T2D ^37^ and obesity associated genes ^4,38–45^. We identified a total of 11 obesity associated genes showing enriched expression in hypothalamic neurons, including *ACHE, ERBB4*, *GRID1*, *KCNK3*, *KCNK9*, *KSR2, NRXN3*, *NTRK2*, *PLXNA4*, *POMC* and *STX1B* (Fig. 4E). Additionally, we identified 13 T2D associated genes out of 257 from the T2D knowledge portal database ^37^ that were enriched in hypothalamic neurons namely *ACVR1C*, *ADCY5, DGKB, DGKG*, *GLP1R, GNG4, POMC, RASGRP1, SCN3A, HCN1, ITPKA, KSR2,* and *NRXN3* (Fig. 4E and 4F). In conclusion, our hPSC-derived hypothalamic neurons exhibit dynamic transcriptome changes that align with prenatal hypothalamus development and show enrichment of genes associated with obesity and T2D, highlighting their relevance and potential utility for modeling metabolic disorders.

### Integrated Analysis of Transcriptomic and Chromatin Accessibility Changes Over the Course of Hypothalamic Neuron Differentiation

To provide insight into the epigenetic modulation occurring over the course of human hypothalamus development, we conducted an assay for transposase-accessible chromatin with sequencing (ATAC-Seq). We profiled a total of 43 samples, aligning with the time-course sample profile used in our transcriptomic analysis (Fig. 3 and Fig. 4). PCA of ATAC-seq samples demonstrated clustering of Day 0, Day 7, and Day 14 samples according to differentiation time points, while samples from Day 21 and Day 28 displayed intermixing (Fig. 5A). This similarity between Day 21 and Day 28 samples corroborated previously observed transcriptomic similarities between the samples from these time points (Fig. 4A and 4C). To better understand how the chromatin landscape changes throughout differentiation and to compare it to transcriptomic changes, we compared the absolute numbers of differentially regulated genes between adjacent differentiation time points (Fig. 5B) and observed a similar trend in ATAC peaks observed in our analysis (Fig. 5C). Volcano plot analysis of differential ATAC-seq peaks between the differentiation start point (Day 0; hPSC) and endpoint (Day 28; hypothalamic neurons) demonstrated significant changes in the chromatin landscape (Fig. 5D). As expected, these ATAC-seq peaks were highly enriched around the transcription start sites (TSS) of genes. Among the top differential peaks with increased or decreased accessibility in hypothalamic neurons were genes associated with obesity and T2D (Fig. 5D and Supplementary table 3).

**Figure 5.**
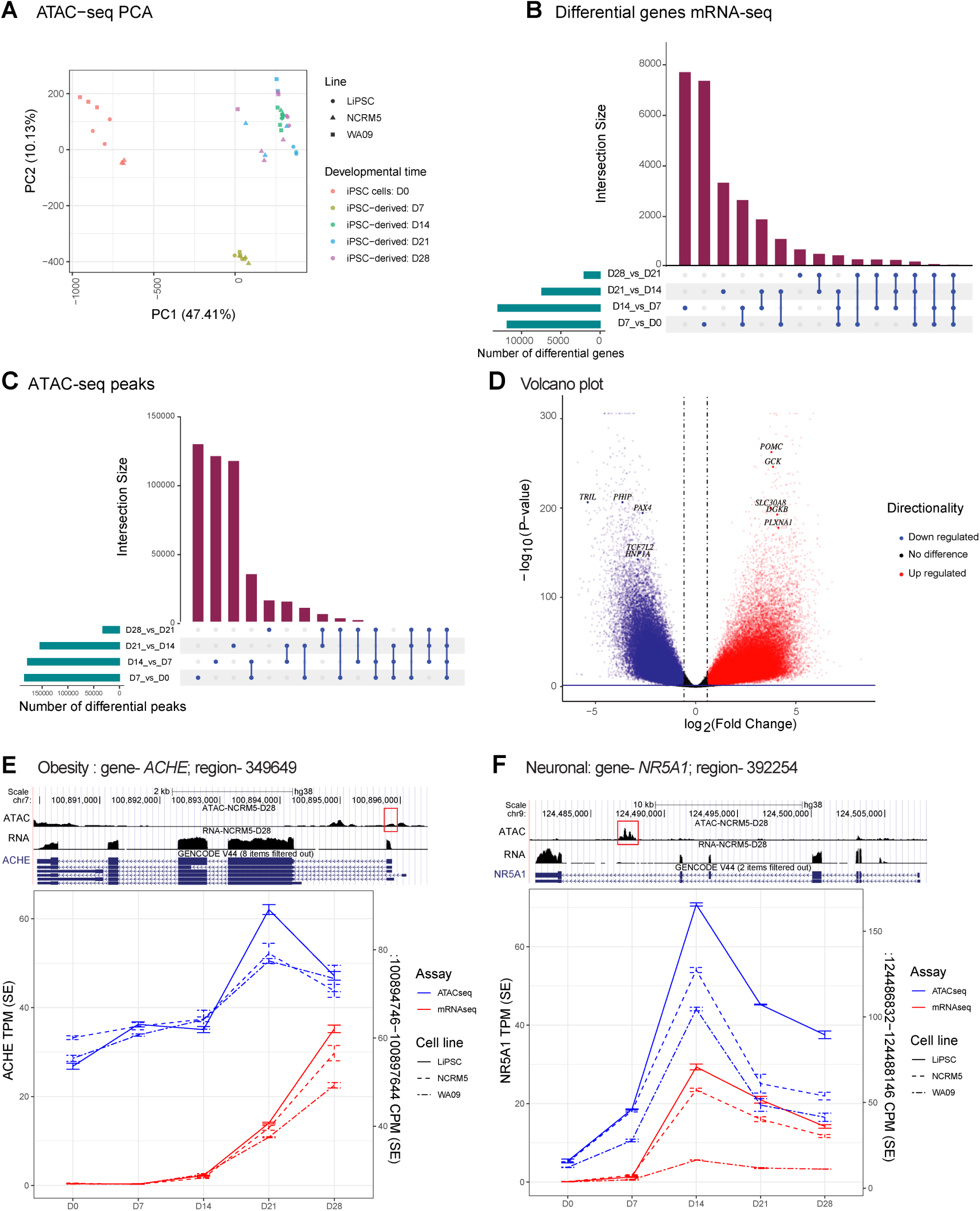
Epigenetic Dynamics of Human Hypothalamus Development Revealed Through ATAC-seq Profiling. (A) PC 1 and 2 of hPSC derived samples from different timepoints of differentiation based on ATAC-seq data. (B) Differential genes shared across comparisons of adjacent time points taking into account directionality of expression changes. Note, by D7 and D14, the largest number of genes are up/down regulated. (C) Similar to Figure 5B, differential peaks shared across comparisons of adjacent developmental time points. Very similar pattern as we observe in the gene expression data. (D) Volcano plot of differential ATAC-seq peaks between D0 and D28. Highlighted are T2D and obesity related genes. (E) *ACHE* association with a potential regulator element in a likely promoter. (F) *NR5A1* association with an intronic ATAC-peak, likely an enhancer. NR5A is an important transcription factor in hypothalamus development.

Next, we sought to identify associations between differential gene regulation and ATAC-peaks detected at intronic or promoter regions, thereby identifying potential regulatory regions or enhancers for the specific gene, as previously described ^46^. We fit a general linear model to assess the association considering the hPSC lines and differentiation timepoint. We detected over sixty thousand accessible chromatin regions associated with gene expression across the differentiation time course (FDR < 0.05; Supplementary Table 3). These include many genes that were previously associated with metabolic disorders and hypothalamus development. Among these, a 2878 base pair (bp) peak around the promoter of ACHE positively correlated with *ACHE* gene expression (Fig. 5E). Likewise, a 1314 bp peak in the last intron of the *NR5A1* gene positively correlated with the time-course expression of this key regulator of medio-basal hypothalamus development ^47^ (Fig. 5F).

In conclusion, our ATAC-seq analysis revealed significant changes in chromatin landscape during the differentiation of hPSCs into hypothalamic neurons, aligning with transcriptomic changes and identifying potential regulatory regions that merit further investigation.

### Functional Characterization of Hypothalamic Neurons

One of the notable impacts of insulin on the brain is its role in regulating feeding behavior. POMC neurons of the arcuate nucleus project into the median eminence outside the blood-brain barrier where they are able to detect circulating levels of insulin, glucose and leptin for appetite regulation ^48,49,50^. Following a meal, circulating insulin levels increase and exert depolarization and activation of anorexigenic POMC neurons, partly mediated by the activation of TRPC-5 channels ^51^. Since complete insulin withdrawal results in massive cell death of neurons ^52^, we developed an assay to assess the functional response of the differentiated hypothalamic neurons to insulin. First, we evaluated their basal spontaneous action potentials using multi-electrode arrays (MEAs) following maintenance in Arc-3 medium with reduced insulin for 96 h (Fig. 6A). After 96 h in reduced insulin conditions, the hypothalamic neurons were then treated with increasing doses of human recombinant insulin (0, 0.01, 0.1, and 1 mg/ml) for two hours and their spontaneous activity was re-evaluated using MEAs. A dose-dependent functional response to insulin was observed via increased mean firing rates of hypothalamic neurons (Fig. 6B). To validate these findings across biological replicates, we treated hypothalamic neurons derived from three independent cell lines to low (0.01 mg/ml) and high (1 mg/ml) insulin for two hours. We observed differential responses in all three lines to increasing insulin concentrations, as measured by changes in mean firing rate (Fig. 6C) and average burst frequency (Fig. 6D).

**Figure 6.**
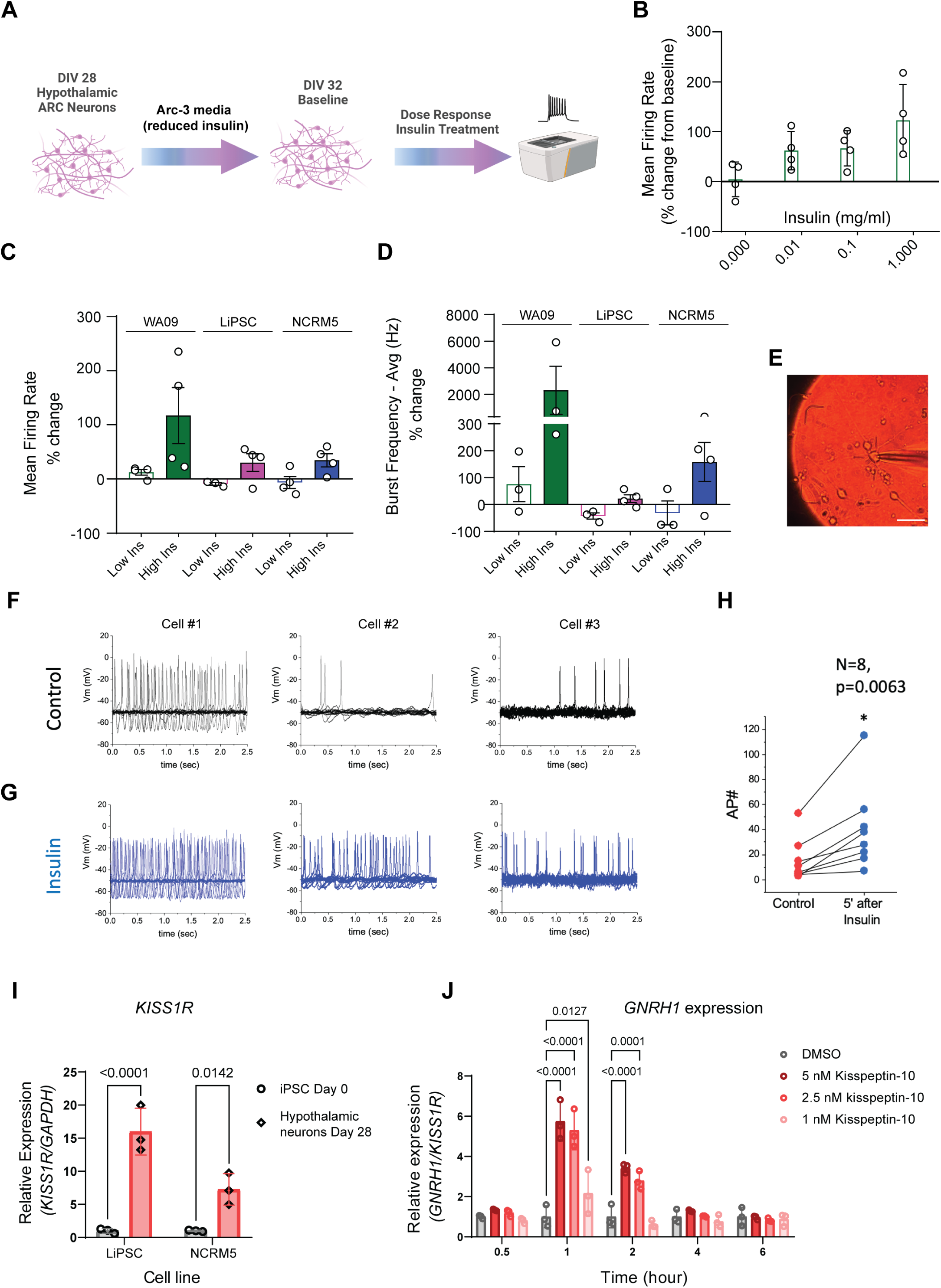
Functional Characterization of Hypothalamic Arcuate Neurons. (A) Schematic overview of MEA assay design for capturing insulin responses. (B) Mean firing rate of hypothalamic arcuate neuron cultures captured after 2 h treatment with increasing doses of human recombinant insulin. Data was normalized to baseline (0 h, no treatment). (C-D) Differential response to low and high insulin concentrations validated across neurons derived from three independent hPSC lines (LiPSC-GR1.1, WA09, NCRM-5) as measured by changes in mean firing rate (C) and burst frequency (D) after 2 h treatment. (E) Representative microscopic image of an arcuate neuron pierced by an electrode for patch-clamp electrophysiology recordings. (F) Spontaneous action potential firing recorded from a holding potential of –50 mV before insulin addition. Three representative neurons derived from the LiPSC-GR1.1 cell line are represented. (G) Spontaneous action potential firing recorded from a holding potential of –50 mV three to five minutes after insulin (0.01 mg/ml) addition. Three representative neurons derived from the LiPSC-GR1.1 cell line are represented. (H) Quantification of action potentials before and after insulin treatment. Note the significant increase in firing rate of insulin-treated cells. N=8, Paired Student’s t test, p=0.0063. (I) KISS1R expression was significantly upregulated in DIV28 hypothalamic neurons as compared to DIV 0 hPSC. Data points represent three replicate samples for the LiPSC-GR1.1 and NCRM-5 lines. (J) Mid-throughput (96-well plate format) RT-qPCR assay for *GNRH1* mRNA expression in response to increasing doses of Kisspeptin-10 (1 nM, 2.5 nM, and 5 nM) or no treatment control. Note the peak increase in *GNRH1* expression at 1 h and its downregulation at later time points captured. Target gene expression level was normalized to that of the housekeeping gene *GAPDH*. Three replicate wells of neurons derived from LiPSC-GR 1.1 were used per treatment group.

To delve deeper into the electrical properties of hPSC-derived hypothalamic neurons, we conducted patch-clamp electrophysiological recordings for eight neurons and demonstrated that patched neurons did not exhibit burst firing but displayed a spontaneous tonic firing pattern at - 50mV (Fig. 6 E-F). When challenged with insulin, there was a noticeable and significant increase in the firing rate of neurons (Fig. 6G and Fig. 6H).

As noted earlier, differential gene expression analysis suggested the presence of a GnRH neuron population at later stages of differentiation (Fig. S5B), and thus we explored this further with both molecular and functional assays. GnRH neurons are a poorly understood subgroup of ARC cells that exhibit pulsatile GnRH secretion in response to neuropeptide kisspeptin to promote release of gonadotropin hormones by the pituitary. In all mammals, this mechanism is essential for both fertility and the onset of puberty ^53,54^. We conducted RT-qPCR to validate *KISS1R* expression in hypothalamic neurons as KISS1R is required for the neuronal response to kisspeptin in the ARC. Utilizing two independent iPSC lines, we observed higher levels of *KISS1R* mRNA in hypothalamic neurons compared to iPSCs (Fig. 6I). Next, we measured the level of *GNRH1* mRNA at various time points (0.5, 1, 2, 4, and 6 h) during a dose-response experiment with escalating dosages (0, 1, 2.5, and 5 nM) of the KISS1R agonist, Kisspeptin-10. *GNRH1* mRNA levels in hypothalamic neurons increased with increasing dosages of Kisspeptin-10 (Fig. 6J). Expression of *GNRH1* peaked 1 h after treatment and returned to baseline 4h later (Fig. 6J). This transient upregulation of *GNRH1* indicates the neurons’ ability to dynamically respond to Kisspeptin-10 pulses and lays the groundwork for the use of this iPSC model in the reproductive endocrinology research.

### Elucidating glucose, insulin and incretin mimetic effects using human hypothalamic neurons

To further demonstrate the translational potential of the hPSC-derived hypothalamic neurons, we assessed their molecular responses to challenges with glucose, insulin, and commercially available incretin mimetic drugs (glucagon-like peptide 1 receptor [GLP1R] agonists). The ARC is well-positioned to react quickly to circulating levels of hormones and nutrients because of its proximity to the portal blood capillary network and the median eminence. Signals of nutrient-state, such as insulin and glucose are important regulators of appetite and satiety ^5^. Therefore, we sought to investigate the potential of our *in vitro* derived hypothalamic arcuate neurons to respond to these hormonal and humoral stimuli. We profiled POMC and insulin receptor (IR, alpha and beta subunits) protein levels following a four-day (96 h) treatment with different media formulations. The experimental conditions included: i. low glucose with insulin; ii. low glucose with reduced insulin; iii. high glucose with insulin; iv. high glucose with reduced insulin; and v. regular Arc-3 medium control. Interestingly, under high glucose conditions, we observed higher levels of POMC proteins compared to those in low glucose conditions (Fig. 7A-B). The strongest increase was observed in neurons exposed to high glucose with reduced insulin (+g/-i), with a two-fold increase compared to low glucose with reduced insulin (-g/-i) (Fig. 7B). In contrast, IR expression exhibited a different pattern of modulation with higher levels of both IR subunits in the reduced insulin conditions, regardless of glucose conditions (Fig. 7A, 7C, and 7D). Incretin receptor GLP1R is expressed in the arcuate nucleus and has recently garnered significant interest. In particular, GLP1R expressed in POMC neurons of the ARC has been implicated as a mediator of weight loss via effects associated with antidiabetic drugs acting as incretin mimetics ^55^. Building on our ESI analysis, which indicated that *GLP1R* is one of the T2D-associated genes enriched in our hypothalamic neurons on day 28 (Fig. 4F), we first measured *GLP1R* transcript expression using RT-qPCR under different culture conditions: low glucose (1 g/L), medium glucose (4 g/L), or high glucose (9 g/L) supplemented Arc-3 medium for 96 h (Fig. 7E). Interestingly, we observed the highest expression of *GLP1R* under medium glucose, although this was not significantly different from the low glucose condition. The high glucose condition displayed a statistically significant reduction in *GLP1R* expression (Student’s t test; p=0.0042, high vs medium glucose p=0.0042; low vs high glucose p=0.045; Fig. 7E). After we confirmed which culture conditions support the expression of GLP1R, we investigated the pharmacological activation of GLP1R and its resultant phosphorylation of CREB. Hypothalamic neurons from 2 independent hPSC lines were challenged with two FDA-approved medications for glycemic management, Metformin hydrochloride and Exendin-4, for periods of 15, 30, or 60 min. Forskolin, a strong adenylate cyclase activator, was included as a positive control as GLP1R is a G-protein coupled receptor (GPCR) and GPCR activation is followed by adenylate cyclase stimulation prior to CREB phosphorylation ^56^. Forskolin treatment resulted in significant phosphorylation of CREB after 15 min (over a 5-fold increase compared to control) and increased at the 30 min and 60 min time points in both cell lines (Fig. 7F-H). Treatment with Exendin-4 and Metformin led to detectable and robust phosphorylation of CREB in both cell lines, several fold higher than the control levels (red dashed lines) at all time points. In summary, these experiments highlight the translational value of hPSC-derived hypothalamic neurons by demonstrating their ability to functional responsiveness to humoral, hormonal, and pharmacological stimuli, such as glucose, insulin, and incretin mimetic drugs. Taken together, these findings support the suitability of this human-based model for T2D and obesity studies and create new opportunities for research of other metabolic and reproductive disorders.

**Figure 7.**
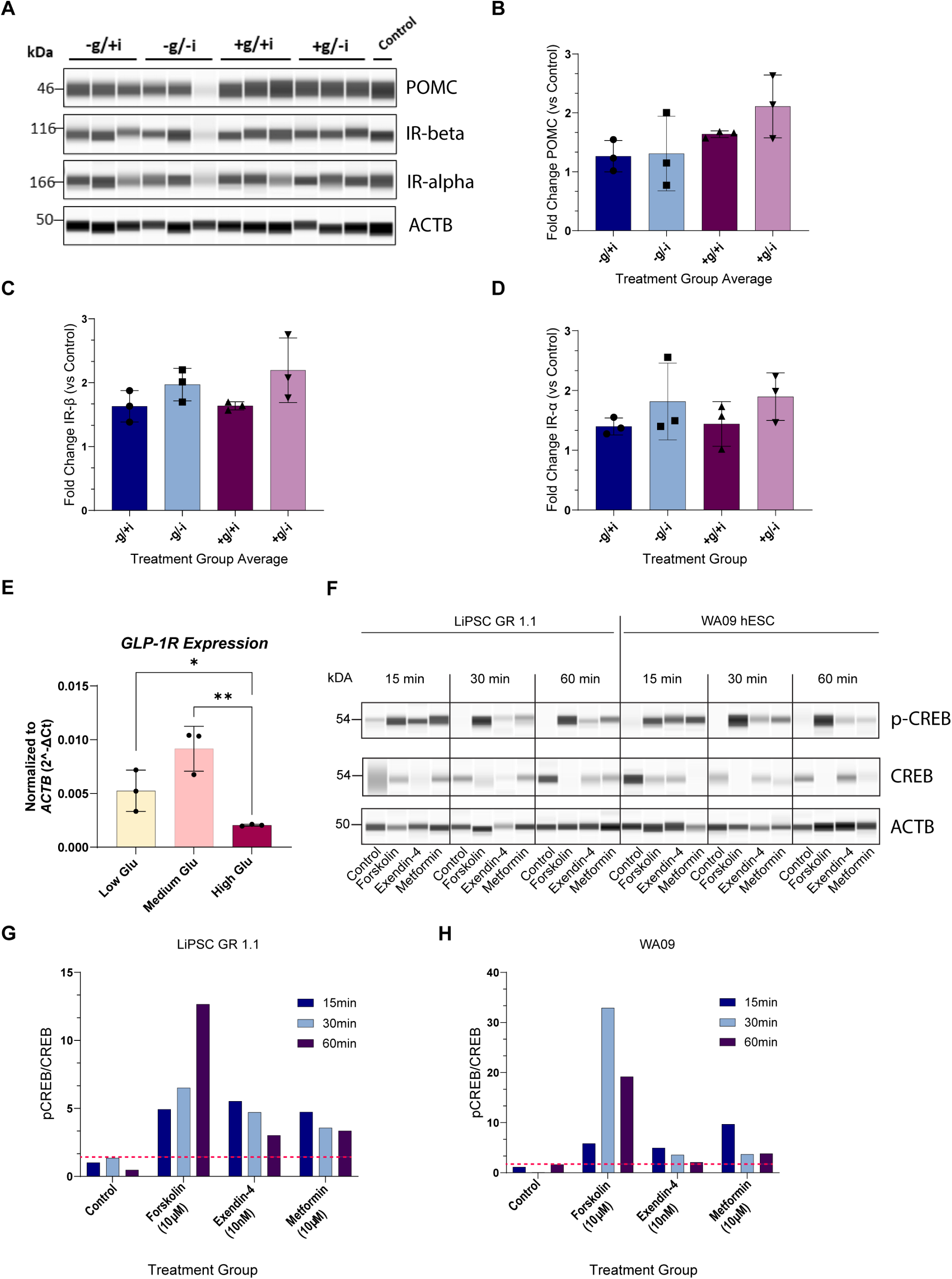
Pulse Assay Experiments to Measure Glucose, Insulin, and Incretin Mimetic Drug Responses. (A) Western blot analysis of triplicate samples of hypothalamic neurons at DIV 28 (LiPSC-GR 1.1) treated with a combination of low (-) or high (+) glucose, with B27 with insulin (+) or with B27 without insulin (-) over a 96 h period. POMC upregulation is observed in high glucose samples. (B-D) Bar graph representations of peak area normalized to Beta Actin (ACTB) and no-treatment control show the strongest expression of POMC in high glucose treated neurons (B), while IR alpha and beta show differential upregulation in samples with reduced insulin compared to those with insulin (C, D). One-way ANOVA F_(3, 8)_ = 2.401; P=0.1432 (B), F_(3, 8)_ = 2.363; P=0.1472 (C), F_(3, 8)_ = 1.066; P=0.4161 (D). (E) RT-qPCR analysis of *GLP1R* transcript expression in iPSC-derived hypothalamic neurons (LiPSC-GR 1.1) treated with Arc-3 medium with low (1 g/L), medium (4 g/L) and high glucose (9 g/L). Unpaired two-tailed Student’s t-test high vs medium glucose **p=0.0042, medium vs low glucose p =0.0753, low vs high glucose *p=0.045 (F) Western blot analysis of CREB phosphorylation in response to GLP1R/GPCR pharmacological activation. Hypothalamic neurons at DIV 28 (LiPSC-GR 1.1 and WA09) were treated for 15, 30, or 60 min with FDA-approved incretin mimetics (Metformin Hydrochloride and Exendin-4), an adenylyl cyclase activator (Forskolin, positive control), and normalized to Beta Actin and no treatment control.

## Discussion

### A Path Towards Translational Research in Neuroendocrinology

The hypothalamus and its arcuate nucleus is a central research focus of neuroendocrinologists studying metabolic disorders ^57–59^ and reproductive dysfunction ^60–63^. Recent advancements in iPSC technology have provided a new avenue for establishing human *in vitro* models of neuropeptidergic hypothalamic neurons ^15–17,22^. While these models have offered valuable insights, they have lacked the scalability required for successful translation to genome-wide genetic and large library drug screens. Primary human hypothalamic neurons are not available to support therapeutic drug discovery, and the use of *ex vivo* human hypothalamus tissue has been limited to single-cell profiling experiments to map hypothalamic cell subpopulations ^10,14^. The human PSC-differentiated hypothalamic neuron model presented in this study consistently achieves a gene expression signature resembling that of the mediobasal hypothalamus across multiple independent hPSC lines. This suggests high reproducibility of the method, which is crucial for its application in translational research, particularly when utilized across various patient-derived iPSCs. Moreover, the strongest rationale for using an iPSC-based model lies in its presumed physiological relevance ^64^. In this regard, the hypothalamic neurons presented here demonstrate responses to pharmacological, hormonal, and humoral stimuli involved in the regulation of appetite and satiety as well as the initiation and maintenance of reproductive competence through the control of gonadotropin release. Given the scalability and reproducibility of the approach, we envision employing these iPSC-derived neurons in comprehensive phenotypic screens. The comprehensive characterization reported in this study, provides compelling evidence that supports their physiological relevance to metabolic disorders such as obesity and T2D, as well as endocrine reproductive disorders.

### hPSC derived ARC Neurons Recapitulate the Prenatal ARC *in vivo*

Our protocol produces a robust population of Arc neurons, including 56% POMC-positive, 15% TH-positive and 6% SST-positive. Our hPSC-derived neurons reach a developmental stage that is comparable to the prenatal hypothalamus within a four-week timeframe. This finding is consistent with other hPSC differentiation protocols that generate neurons with characteristics of a prenatal stage in brain development ^65,66^. In line with recent single cell profiling studies of human hypothalamic development, appetite-promoting populations such as AgRP/NPY neurons were not observed in our cultures, as they emerge during later stages of development ^14^. Potential promoter regulatory elements and gene enhancer regions identified in this study can serve as foundation for further functional validation studies. Subsequent studies can also explore approaches to induce epigenetic modifications to further enhance the maturity of the derived cells ^67,68^. ARC dopamine neurons, expressing TH and GABA, tonically inhibit prolactin release from the pituitary and can be modulated by endogenous opioids met-enkephalin and dynorphin ^12,13,69^. These neurons comprised 10-15% of the total neurons generated using our approach and provide additional value to the established *in vitro* model of ARC neurocircuitry. However, further studies are needed to establish functional assays targeting dopaminergic subpopulation.

### Resource for Modeling Hypothalamic Pathophysiology in Metabolic Disorders

Understanding the molecular and functional responses of these hPSC-derived neurons to metabolic signals could provide valuable insights into the pathophysiology of metabolic disorders associated with the hypothalamus. The expression of T2D and obesity associated genes ^37,38^ in these hPSC-derived hypothalamic neurons further support the use of this resource in future studies modeling metabolic disorders. Insight into the mechanisms underlying susceptibility to weight gain and insulin resistance due to increased caloric intake is crucial in combating these prevalent metabolic diseases. Previous studies have found that selective decrease in arcuate nucleus insulin receptor levels, and increase in POMC can cause weight gain, increased adiposity, and rapid onset of hepatic insulin resistance ^70,71^. The dynamic modulation of the POMC and IR alpha and beta protein levels in response to glucose and insulin, as demonstrated here, indicate their utility in high-throughput assays and phenotypic screens. Multi-omics datasets presented here should serve as a rich data mining resource and help provide insight into the development of the hypothalamus and arcuate nucleus. Finally, the combination of this robust differentiation method with a fully automated robotic platform will allow for the generation of hundreds of millions of human hypothalamic neurons from various patient lines carrying obesity and T2D associated variants and create opportunities for cross institutional collaborations for the core efforts of the Hypothalamus Consortium, to address metabolic disorders such as obesity and T2D.

## Experimental Procedures

### Cell Culture

hPSC lines were maintained in E8 with vitronectin (VN) as a coating substrate and passaged with the use of CEPT for a minimum of 7 passages before initiating differentiation on coated microplates or T175 flasks.

### Differentiation of hPSCs into Hypothalamic Neurons

For detailed instructions and media compositions, refer to Supplementary Methods. Briefly, hPSCs were detached using 0.5 mM EDTA (ThermoFisher, 15575020). The resulting cell clumps were counted using the Nexcelom Cellometer automated cell counter and plated at a density of 2 x 10^4^ cells/cm^2^ on VN-coated microplates or T175 flasks in E8 medium supplemented with CEPT (day −1). The following day, the medium was replaced with freshly prepared Arc-1. On day 7, cells were dissociated to single cells using Accutase and plated on AggreWell plates to form spheres. These spheres were maintained in Arc-2 medium with daily media changes. Day 11, the media was replaced with Arc-3 medium. On day 14, the spheres were dissociated into single cells using the Miltenyi Biotec embryoid body dissociation kit (130-096-348) and plated on Geltrex-coated microplates at a density of 1×10^5^ cells per cm^2^ in Arc-3 media. Every other day, a half media change was performed until day 28, when most of the molecular and functional assays in this study were conducted.

### Automated Cell Culture

Scalable robotic cell culture and differentiation were carried out using the CompacT SelecT platform as previously described ^19^ and following the steps described above. For sphere formation, T175 flasks were pretreated with anti-adherence solution (STEMCELL Technologies).

### Immunocytochemistry and High Content Imaging Analysis

hPSC-derived hypothalamic neurons at different time points were cultured as described above on glass-bottom multiwell plates (Cellvis, P24-1.5H-N). The cultures were fixed with 4% formaldehyde (Thermo-Fisher, 28908) in PBS for 15 min. Then the cultures were incubated with PBS + 4% donkey serum +0.1% Triton-X for 1 h at room temperature, followed by incubation with primary antibodies overnight at 4°C. Secondary antibodies were incubated at room temperature for 1 h. Refer to Table S1 for complete list of primary and secondary antibodies. The cultures were stained with 4 µM Hoechst 33342 (VWR, PI62249) in PBS for 10 min before imaging on the Opera Phenix high-content microscope (Revvity) or Zeiss LSM 710 confocal microscope.

### RNA seq

iPSCs from two different healthy individuals (LiPSC-GR1.1 and NCRM-5) and hESC (WA09) as well as derived neural progenitors and hypothalamic neurons at different differentiation time points were lysed using RLT plus buffer (Qiagen, 1053393) supplemented with 2-mercaptoethanol (Millipore-Sigma, 63689) directly in wells and RNA was extracted and purified using RNeasy Plus Mini Kit (Qiagen, 74136) according to the manufacturer’s instruction. QIAcube automated workstation was used for the extraction (Qiagen). Genomic DNA was eliminated by both the gDNA eliminator column and on-column incubation with DNase I (Qiagen, 79256). Sequencing libraries were constructed and sequenced at the Azenta sequencing facility using Illumina TruSeq® Stranded mRNA kit.

### ATAC-seq

ATAC-seq was performed on 50.000 to 100.000 cells according to the manufacturer’s instructions (Active Motif, 53150). The cells were pelleted and washed with 100 μL of ice-cold PBS. They were then lysed with 100 μL of ATAC Lysis Buffer on ice for 3 min and centrifuged at 500 g for 10 min at 4°C. The nuclei were incubated in 50 μL of Tagmentation Master Mix at 37°C for 30 min in a thermomixer set to 800 RPM. During this process, the Tn5 Transposase probed accessible DNA regions in the nuclei by inserting sequencing adapters into open regions. The tagged DNA was then purified and PCR-amplified. Prior sequencing, the libraries were quantified by qPCR using the KAPA library quantification kit for Illumina platforms (Roche, KR0405) using QuantStudio^TM^ 12K Flex Real-Time PCR System (Thermo Fisher Scientific). All samples were normalized according to concentration and pooled. Libraries were loaded and sequenced using the NovaSeq 6000 System (Illumina).

### Patch-clamp Electrophysiology

For these experiments, human iPSC-derived ARC neurons were assessed between days 28 and 34 after plating. On the day of recording, the extracellular media was switched from DMEM/F12 media to BrainPhys media (Stem Cell Technologies, 05790). The composition of the intracellular solution was (in mM): 130 K-glutamate, 9 KCl, 10 NaCl, 0.5 MgCl2, 4 Na2ATP, 10 HEPES, pH adjusted to 7.3 with KOH, osmolarity: 292 mOsm/kg. Membrane potential measurements were performed utilizing the whole-cell configuration of the patch-clamp technique in current-clamp mode with a commercial amplifier (HEKA Instruments, EPC-10) and pipettes of borosilicate glass (Warner) with resistances of 5-8 MΩ when filled with the internal solution. Neurons exhibiting signs of qualitative clamp error or with a holding current >1 nA were excluded from the analysis. After establishing the whole-cell current-clamp configuration, we waited for the membrane potential to stabilize for 3 min. Spontaneous action potential firing was monitored for 14 sec from a holding potential of –65 or –50 mV before and 3-5 min after adding insulin to the recording chamber. In some cases, action potentials were initiated by 20-ms suprathreshold depolarizing current pulses. The amplifier was utilized in the low-frequency voltage-clamp mode, a modified current-clamp mode that functions like a current clamp for fast signals and a voltage clamp for low-frequency signals. Electrical signals were typically low-pass filtered at 3-10 kHz (3-pole Bessel filter). The sampling frequency was 10 kHz. Data were acquired, digitized, stored, and analyzed using commercial software (Patchmaster and Fitmaster; HEKA Instruments). For further analysis, Origin-Pro software (OriginLab, Northampton, MA) was employed.

### Multi-electrode array (MEA)

Neuronal activity was analyzed using the Maestro Pro multi-electrode array system (Axion Biosystems) according to the manufacturer’s protocol. Briefly, day 14 hypothalamic neurons were thawed in Arc-3 medium with CEPT and plated at a density of 80,000 cells/well on PLO/Laminin coated 48-well or 96-well MEA plates. Twenty-four hours post-plating the medium was replaced by Arc-3 medium and full media change was performed every 2-3 days. On day 28, to reduce insulin media was changed to Arc-3 with B27 without insulin, and hypothalamic neuron activity was recorded on day 32, 18 days post plating. Human recombinant insulin concentration used were as follows: 0.01 mg/ml (low insulin), 0.1 mg/ml and 1 mg/ml (high insulin).

### Differential Glucose, Insulin, and Incretin Mimetic Treatments

Arc-3 media was modified as follows to create four media conditions: low glucose [1 g/L] and B27 with insulin, low glucose [1 g/L] with B27 without insulin, high glucose [9 g/L] and B27 with insulin, high glucose [9 g/L] with B27 without insulin. Matured hypothalamic cultures were treated with one of the four modified ARC3 conditions for four days, with 3/4th media change on day 29 and day 30. On day 32, cells were gently rinsed with PBS (-/-) two-three times and lysed with RIPA buffer (ThermoFisher Scientific, 89900) supplemented with Halt Protease Inhibitor Cocktail (ThermoFisher Scientific, 78440) and collected. Lysed cells were sonicated, and debris-free lysis was collected for BCA assay and Western Blot Analysis.

Incretin mimetic drug treatments were performed by supplementing unmodified Arc-3 medium with Metformin hydrochloride at 10 µM (Tocris, 2864), Exendin-4 at 10 nM (Tocris, 1933), Forskolin (Tocris, 1099) at 10 µM. Cells were rinsed with PBS (-/-) once then lysed with RIPA buffer supplemented with Halt Protease Inhibitor Cocktail after 15 min, 30 min, 1 h, 2 h, or 24 h dependent on the media treatment. Lysed cells were sonicated, and debris-free lysis was collected for BCA assay and Western Blot Analysis.

### Western Blot

The Jess automated Western blotting systems (ProteinSimple) were utilized for quantitative analysis of protein expression, as previously described ^18^. All Western blot data are presented by lanes in virtual blot-like images. In brief, cells were harvested by scraping, pelleted, washed with PBS, flash-frozen using dry ice, and stored at −20 °C until processed. Cell pellets were then resuspended in RIPA buffer (ThermoFisher Scientific, 89900) supplemented with halt protease and phosphatase inhibitor cocktail (ThermoFisher Scientific, 78440) and lysed by sonication. Lysates were subsequently cleared of debris by centrifugation at 14,000g for 15 min and quantified using the BCA protein assay kit (Thermo Fisher Scientific). Protein quantification was performed using the 12-230 kDa plates (ProteinSimple, PS-MK15) in a JESS Capillary Western Blot analyzer following the manufacturer’s recommendations. Protein quantification was conducted using the Compass software. For a full list of primary and secondary antibodies used, please refer to Table S1.

### Real-Time Reverse Transcription Polymerase Chain Reaction (RT-PCR)

RNA was extracted using the SingleShot Cell Lysis Kit (Bio-Rad Laboratories, 1725080) per the manufacturer’s instructions. Briefly, a 96-well plate was loaded with 100k cells per cm^2^ of iPSC-derived human hypothalamic neurons (LiPSC-GR 1.1) in complete Arc-3 medium. After each treatment, cells were lysed for 10 min at room temperature in 40 μL of lysis buffer containing DNase and proteinase, followed by 5 min at 37°C and 5 min at 75°C. Each sample underwent RT-PCR in triplicate using Reliance One-Step Multiplex Supermix (Bio-Rad Laboratories, 12010220) and specific probe sets for each gene. The total volume of each reaction was 10 μL, and the final concentration of each gene’s primer is 1×. On a QuantStudio^TM^ 3 Real-Time PCR System, the real-time PCR reactions were performed according to the manufacturer’s instructions (1 cycle of 50°C 10 min and 95°C 10 min for reverse transcription and enzyme activation, followed by 40 cycles of 95°C 15 s and 60°C 30 s for annealing and extension). Primer probe sets *KISS1R* (Hs00261399_m1), *GNRH1* (Hs00171272_m1), and *GAPDH* (4310884E) were all purchased from ThermoFisher Scientific. Using the comparative threshold cycle (Ct) method, the target gene’s expression level was normalized to that of the housekeeping gene *GAPDH*. Kisspeptin-10 was purchased from TOCRIS (TOCRIS, 2570). For *GLP1R* RT-PCR experiments, differentiated neurons (LiPSC-GR 1.1) at day 32 were cultured for 96 h in one of the three different glucose conditions (low glucose 1 g/l, medium 4 g/L and high 9 g/L), followed by RNA isolation using the RNeasy Plus 20 mini kit (Qiagen, 74136) in the QIAcube equipment. cDNA was synthesized using the high-capacity RNA to cDNA kit (Applied Biosystems, 4387406). PrimeTime Gene Expression Master Mix (IDT, 1055771) was used to perform the qRT-PCR in the QuantStudio 12K Flex Real Time PCR system. *GLP1R* (Hs.PT.58.39163702.g) and *ACTB* primers (Hs.PT.39a.22214847) were purchased from IDT.

### RNA-seq read alignment, quantification, and quality control

We processed RNA-seq reads as described in Xue et al. ^46^. Briefly, we aligned the sequencing reads to the GRCh38 genome assembly using STAR (v2.73a) ^72^ with default parameters and quantified expression levels of Gencode v19 genes (Ensembl release 103) using QoRTs (v1.3.6) ^73^ per library/replicate. We generated a raw read count matrix of gene by library and a normalized mRNA expression matrix of transcripts per million (TPM). On average, we generated 60,111,933 (49,044,888 - 70,752,140) paired-end reads per library, of which 89.95% were uniquely aligned to the genome. Out of the aligned reads, 48.33% were unambiguously assigned to unique genes, indicating high quality data.

We normalized the gene by library raw count matrix to TPM considering transcript length and library size information. We excluded genes with no reads in at least one library and retained only protein-coding genes. We scaled the TPM data using the “scale” function of R package (version 3.5.1) and performed a Principal Component Analysis (PCA) using the PCA function of R and used ggplot2 (version 3.5.1) for generating figures. We did not find any outlier replicate per iPSC line at any time point during differentiation. We used a similar normalization approach to compare the transcriptome of iPSC derived cells with data obtained from previously published studies.

### Differential gene expression analysis

We performed differential gene expression analysis as described in Xue et al. ^46^. We assessed gene expression changes in the iPSC lines at each differentiation stage (D7, 14, 21, or 28) relative to the initial iPSC stage (D0), as well as that of adjacent differentiation stages (D0 vs D7, D7 vs 14,…) using R package DESeq2 (v1.42.1) ^74^. We tested genes with read Counts Per Million (CPM) > 0.5 in at least 50% libraries for each comparison. We normalized the raw read counts using the median-of-ratios method, which accounts for differences in library size and RNA composition between samples. A generalized linear model was fit to the count data and the Wald test was used to identify differentially expressed genes. P-values were adjusted for multiple testing using the Benjamini-Hochberg (BH) procedure to control the false discovery rate (FDR). Genes with an FDR < 5% and a fold change (FC) > 1.5 were considered significantly differentially expressed. Gene set enrichment analysis To assess the impact of changes in gene expression through differentiation, we selected the top differentially expressed genes (FDR < 10e-6 and FC >=4) and evaluated the representation of these genes in previously known biological pathways of KEGG ^75^ using the “enrichKEGG” function of the clusterProfiler (v4.11.0) R package ^76^. P-values were adjusted for multiple testing using the Benjamini-Hochberg (BH) procedure to control the FDR, and gene sets with FDR < 5% were considered to be over-represented in the differential genes.

### Gene Expression Specificity Index

We normalized the raw read counts by library size and transcript length for each gene and generated a TPM matrix of differentiation time point by gene. We applied CELLEX v1.2.2 ^35^ on the TPM matrix and calculated expression specificity index for each gene at a given time point.

### Gene Set enrichment and RUV-seq analysis

Gene set enrichments and pathways analysis were performed using Enrichr on the top 100 differentially expressed genes by fold-change, using the KEGG 2021 Human, GTEx Tissues (v8 2023), and Human Gene Atlas databases. Heatmaps were generated using the pheatmap package in R (version 1.0.12), and boxplots were generated with ggplot (release 3.4.1) using transcripts per million abundances calculated by DESeq2.

For comparison to the Rajamani, et al. study ^15^, sequencing data was downloaded from SRA ^77^ Bioproject PRJNA376498. Samples from control iPSC-derived hypothalamic neurons or adult hypothalamic tissue were used for the analysis. The resulting datasets were normalized using upper-quartile normalization ^78^. As no spike-in controls were available, a first-pass analysis was performed using edgeR with a negative binomial GLM, and all genes but the top 5000 by edgeR p-value were used as empirical control genes to normalize all samples together using RUVseq RUVg (version 1.36.0). For comparison to arcuate nucleus neurons, a set of marker genes was derived from the Zhou, et al. publication ^10^, and normalized count values, using the offsets derived from RUVg were used.

### ATAC-seq read alignment, quantification, and quality control

We processed ATAC-seq reads and quantified sequencing reads using the approach used in Xue et al. ^46^. Briefly, we removed duplicate reads and retained 97,076,117 uniquely aligned primary reads per library/replicate on average (minimum of 41,330,939) for downstream analyses. Using the filtered reads, we called ATAC peaks as described in Rai et al. ^79^, removing candidate peaks that overlap with ENCODE blacklists ^80^ and controlling for a FDR of 5%. For each iPSC line at a differentiation time point (D0, 7, 14, 21, and 28), we merged peaks across replicates and retained peaks present in ≥2 replicates. Next, we created a master set of peaks by merging peaks across all libraries, generating a total 395,573 peaks. Finally, we used this master set of peaks to quantify the number of reads mapping to the peaks within each library and generated an accessible chromatin region count matrix. We used MultiQC ^81^ to generate and aggregate QC metrics across libraries and determined all libraries are of good quality (both “per_base_sequence_quality_scores” and “per_sequence_quality_scores” meeting thresholds of “pass”). We applied ataqv (v1.2.1) ^82^ to generate QC metrics, such as the number of high-quality autosomal alignments (HQAA) and transcription start site (TSS) enrichment. We identified no outlier library.

### Differential chromatin accessibility analysis

We performed differential chromatin accessibility analysis using the ATAC-seq read count matrix corresponding to the comparisons in differential gene expression analysis using the approach used in Xue et al. ^46^. We considered peaks with FDR < 5 and a FC > 1.5 to be significantly different open chromatin regions.

### Association of gene expression and chromatin accessibility

We annotated any ATAC-seq peak with a gene where the ATAC-seq peak overlaps 50kb flanking regions of the TSS or gene body (gencode v43, https://www.gencodegenes.org/human/). To infer association of likely cis-regulatory elements and nearby gene expression, we fit a general linear regression to model the inverse-normalized TPM for a gene as the dependent variable, the inverse-normalized ATAC-seq peak read counts as the independent variable taking into account the iPSC line and differentiation time point (0, 7, 14, 21 and 28 day) as covariables (“lm” function from the R package stats (v4.3.2)). Since RNA-seq and ATAC-seq libraries of each iPSC line were not simultaneously generated, we summed up raw counts of three replicates for each gene/peak of a cell line, calculated CPM prior to the inverse normalization. We used the Benjamini-Hochberg procedure to control for the number of tests and considered tests with FDR<5% and FC > 1.5 to be associated.

### Data Availability

Data are presented as the mean ± SD. Statistical analyses (GraphPad Prism) were performed using different tests as appropriate and as described in the figure legends. Bulk RNA-seq and ATAC-seq

FASTQ files have been deposited into the Sequence Read Archive (SRA) under BioProject PRJNA1085406.

## Author Contributions

V.M.J., C.A.T., C.D., F.S.C., S.C. conceived the study and experiments. V.M.J., K.M., C.S., C.D., S.Y., S.R., E.HO., D.F.B., G.RS. and D.B. performed all experiments. T.R., N.N., T.Y., N.S., H.J.G., J.I. performed the bioinformatics analyses. V.M.J., C.L., M.X., M.S., E.HO., M.E., L.B., S.C., D.C., A.S., F.S.C. and C.A.T. contributed to data analysis and discussions. V.M.J. wrote the manuscript with contributions from C.A.T., C.A.D., F.S.C., L.B., S.C. and H.J.G.

## Lead Contact

Further information and requests for resources and reagents should be directed to and will be fulfilled by the lead contacts Vukasin Jovanovic and Carlos Tristan.

## Acknowledgments

Authors would like to thank Matthew Hall, Ann Knebel, Hannah Baskir, Yeliz Gedik, Glib Lirazan, Kathryn Garish, Christopher Cherry and Charles “Pepper” Bonney for their support throughout this work and Ilyas Singeç for his oversight of conceptualization and development of the protocol. We are grateful to Allan Hoofring and Erina He from the NIH Medical Arts Design Section for art designs. We also gratefully acknowledge funding from the Regenerative Medicine Program (RMP) of the NIH Common Fund and by the intramural research program of the National Center for Advancing Translational Sciences (NCATS), and the Interagency Agreement #NTR 12003 from the National Institute of Environmental Health Sciences (NIEHS)/Division of Translational Toxicology (DTT) to the NCATS, NIH. C.A.D. is supported by the Frontiers program of the Russell Berrie Foundation, New York Obesity Research Center, P30 DK026687-43 and DK52431-25. The funders had no role in study design, data collection, and analysis; decision to publish; or preparation of the manuscript. Figure 1A and 2A were generated using Biorender.com.

