## Supplementary materials for "Scalable Hypothalamic Arcuate Neuron Differentiation from Human Pluripotent Stem Cells Suitable for Modeling Metabolic and Reproductive Disorders"

### Supplementary Experimental Procedures

#### Method for Scalable Differentiation of hPSC into Hypothalamic Arcuate Neurons

The hPSCs utilized in this study underwent mycoplasma testing prior to differentiation and karyotype analysis to exclude common cell culture contamination and chromosomal abnormalities. Each cell line was maintained for a series of 7 passages under feeder-free conditions using E8 Medium (ThermoFisher, A1517001) on vitronectin (ThermoFisher, A14700) as a coating substrate. CEPT [1] was included at each passage for 24 h to support viability.

On Day (-1), 20,000 hPS cells/cm$^2$ were seeded on vitronectin-coated vessels in E8 +CEPT (e.g., 180,000 cells per well of a 6-well plate or 3.5 million cells per T175 flask).

On day 0, the media was replaced with Arc-1 medium. From Day 0 to Day 6, full media changes were performed daily, without passaging.

Arc-1 medium composition:

- E6 medium (ThermoFisher, A1516401)
- 100 nM LDN189193 (Tocris, 6053)
- 2 µM A83-01 (Tocris, 2939)
- 4 µM XAV939 (Tocris, 3748)
- 0.5 µM SAG21K (Tocris, 5282)
- Jagged-1 (1277-JG), LIF (7734-LF), CNTF (257-NT), Delta-like protein – 1 (DLL1) (R&D systems), all at 10 ng/ml.

On day 7, cells were dissociated with StemPro Accutase (ThermoFisher, A1110501) for a 5-minute treatment at 37°C (1 ml per well of a 6-well plate; 10 ml for T175 flask), and a cell suspension of 2.5 million cells per ml was prepared in Arc-2 medium with CEPT.

Arc-2 medium composition:

- DMEM/F12 (ThermoFisher, 11320033) + N2 (ThermoFisher, A1370701) + B27 w/o vit. A (ThermoFisher, 12587010)
- 4 µM XAV939 (Tocris, 3748)
- 25 ng/ml Activin A (R&D systems, 338-AC)
- 20 ng/ml BDNF (R&D systems, 11166-BD)
- 20 ng/ml GDNF (R&D systems, 212-GD)
- 200 µM Ascorbic Acid (Tocris, 4055)
- 50 µM db-CAMP (Tocris, 1141)

For ArcSphere formation, 3 ml of prepared cell suspension was added per well of an AggreWell 800 6-well plate (StemCell Technologies) to achieve 5000 cells per microwell, and plates were maintained in a cell culture incubator overnight. The next day, spheres were carefully transferred using a 1 ml wide bore pipet into 40 ml of Arc-2 medium and placed in T175 flasks pre-rinsed with an anti-adherence solution (StemCell Technologies, 07010). One full AggreWell plate was transferred into each T175 flask.

Spheres were maintained in Arc-2 medium until day 11, with daily medium changes. On day 11, the medium was replaced with Arc-3 medium.

Arc-3 medium composition:

- DMEM/F12 + N2 + B27 complete (ThermoFisher, 17504044)
- 20 ng/ml BDNF (R&D systems, 11166-BD)
- 20 ng/ml GDNF (R&D systems, 212-GD)
- 200 µM Ascorbic Acid (Tocris, 4055)
- 50 µM db-CAMP (Tocris, 1141)
- 1 µM DBZ (Tocris, 4489)

On day 14, spheres were dissociated using the neural tissue enzyme dissociation kit and platform (Miltenyi Biotech). At this point, cells were cryopreserved or plated for final maturation.

Cryopreservation medium: Neuron freezing medium (Cell Applications, 042-50) with CEPT.

For final maturation, 100,000 dissociated cells were seeded per $cm^2$ of Geltrex (ThermoFisher, A1569601) coated vessels in Arc-3 medium with CEPT. Every other day, half-media changes were performed until day 28. On day 28, cells can be used for functional assays described in this study.

### Supplementary Figure Legends

**Figure S1. Automated differentiation of hPSCs into hypothalamic neurons**

(A) Schematic overview of the automated differentiation process using the Compact SelecT robotic cell culture system with T175 flasks. Note that the cells are cryopreservable at day 14. Starting with 3.5 million hPSCs per flask and carrying only 3 flasks during the sphere stage, the automated differentiation results in approximately 100 million hypothalamic neurons.

(B) Hypothalamic neurons are available "off the shelf" and require thawing in Arc-3 media with CEPT and 14 days of maturation before assaying.

**Figure S2. Expression of POMC, GABA, SST, and Calbindin in ARC Neuron Cultures**

(A) Co-immunostaining for POMC (red), SST (white), and GABA (green).

(B) Co-immunostaining for Calbindin (d28k; red) and GABA (green).

**Figure S3. Transcriptomic comparison of hypothalamic neurons to publicly available datasets**

(A, B) Gene pathway enrichment analysis using the top 100 genes upregulated at day 28 vs day 0 ($log_2$FC) results in the hypothalamus as the main hit in GTEx and Human Gene Atlas human tissue databases.

(C) Gene pathway enrichment analysis using the top 100 genes upregulated at day 28 vs day 0 ($log_2$FC) results in circadian entrainment as the top hit in the KEGG pathway analysis.

(D) PCA plot showing clustering of samples from this study as compared to samples from Rajamani et al.'s 2018 study. Note the separation in clustering of the samples between the studies.

(E) Heatmap of differential gene expression for a set of genes shown to be markers of 6 major neuronal populations in the human arcuate nucleus adapted from Zhou et al.[2] Note the presence of a large cluster in the middle, resulting from samples of this study (black box).

**Figure S4. Time-course analysis of neural progenitor marker expression and markers of other hypothalamic nuclei.**

(A) Schematic representation of hypothalamic development and key transcription factors involved in hypothalamic nuclei specification. Adapted from Xie and Dorsky [3].

(B) Expression of neural progenitor marker genes NES, SOX2, FABP7, and proliferative marker MKI67 over the differentiation time course.

(C) Time-course expression of arcuate (ARC) nucleus gene markers *ISL1, POMC, NKX2-1, SIX3, AGRP*.

(D) Time-course expression of dorsomedial hypothalamus (DMH) marker *GPR50* and ventromedial hypothalamus (VMH) marker *NPTX2*.

(E) Time-course expression of mammillary nucleus (MN) and suprachiasmatic nucleus (SCN) markers *LHX1, FOXB1*, and *RGS16*.

(F) Time-course expression of zona incerta (ZI) marker *MEIS2*, supramammillary nucleus (SMN) marker *BARHL1*, lateral hypothalamus (LH) marker *CARTPT*, and paraventricular nucleus of the hypothalamus (PVH) markers *CRH* and *OXT*.

Gene markers representing different hypothalamic nuclei are adapted from Herb et al [4].

**Figure S5. Pathways enriched with up-regulated and down-regulated genes across the developmental time points.**

(A-C) Pathways enriched (FDR<0.05) at each time point with up-regulated genes at FDR<1e-6 and |fold change| > 4 compared to Day 0. Rich factor = the number of up-regulated/down-regulated genes overlapping with the respective pathway to the total number of genes in the pathway for a comparison.

(D-F) Pathways enriched (FDR<0.05) with up-regulated genes (E) or down-regulated genes (F) FDR<1e-6 and |fold change| > 4) compared to a previous time point (adjacent time point comparison).

**Supplementary Table 1.** List of primary and secondary antibodies used in this study. Note that Western blot experiments were performed using the automated Jess system (ProteinSimple).

**Supplementary Table 2.** List of genes associated with the analysis presented in Figure 4D.

**Supplementary Table 3.** List of ATAC-seq peaks significantly associated with gene expression over the time course.
