## Supplementary figures and images for "Scalable Hypothalamic Arcuate Neuron Differentiation from Human Pluripotent Stem Cells Suitable for Modeling Metabolic and Reproductive Disorders"

### Supplementary figures

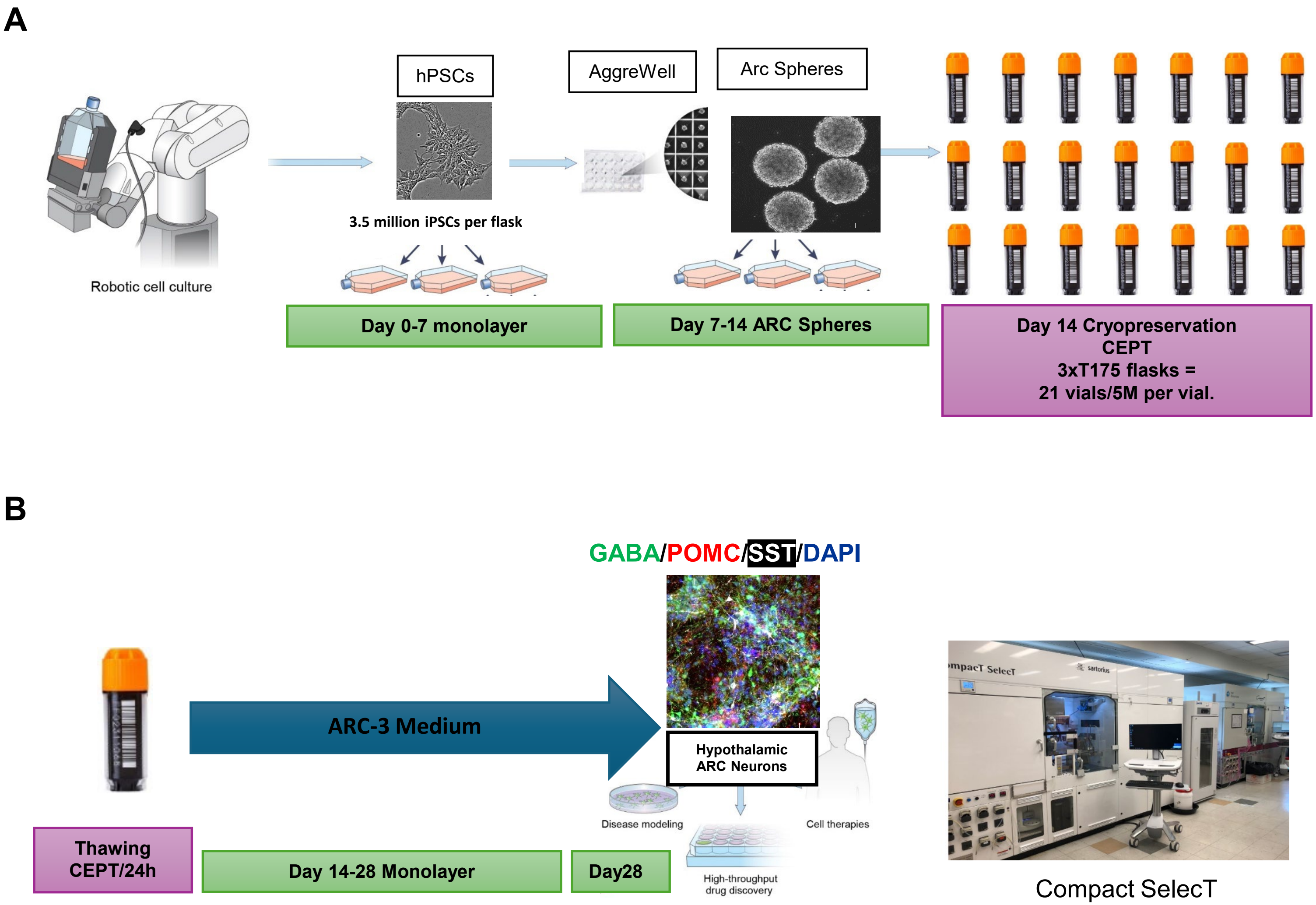

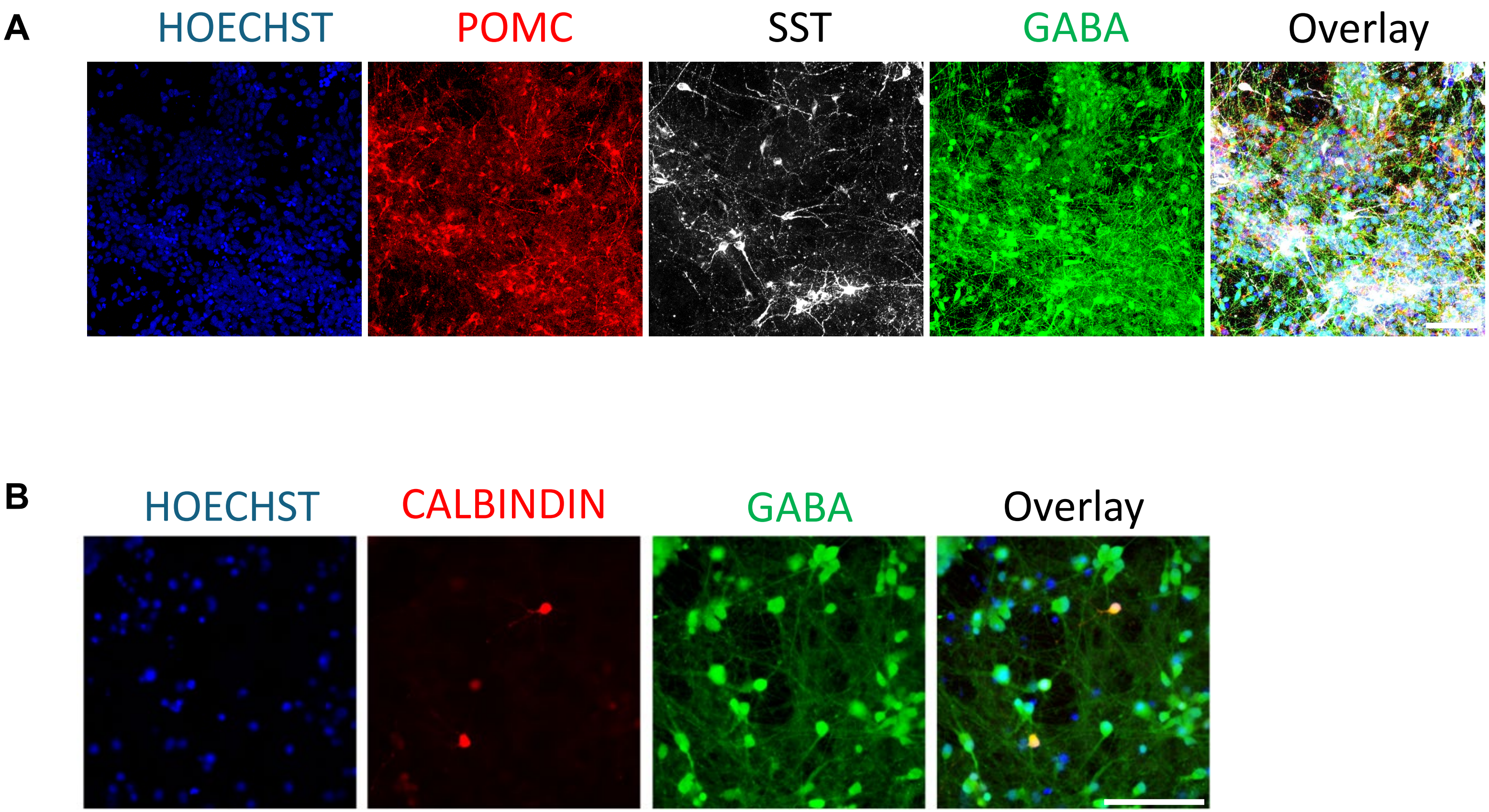

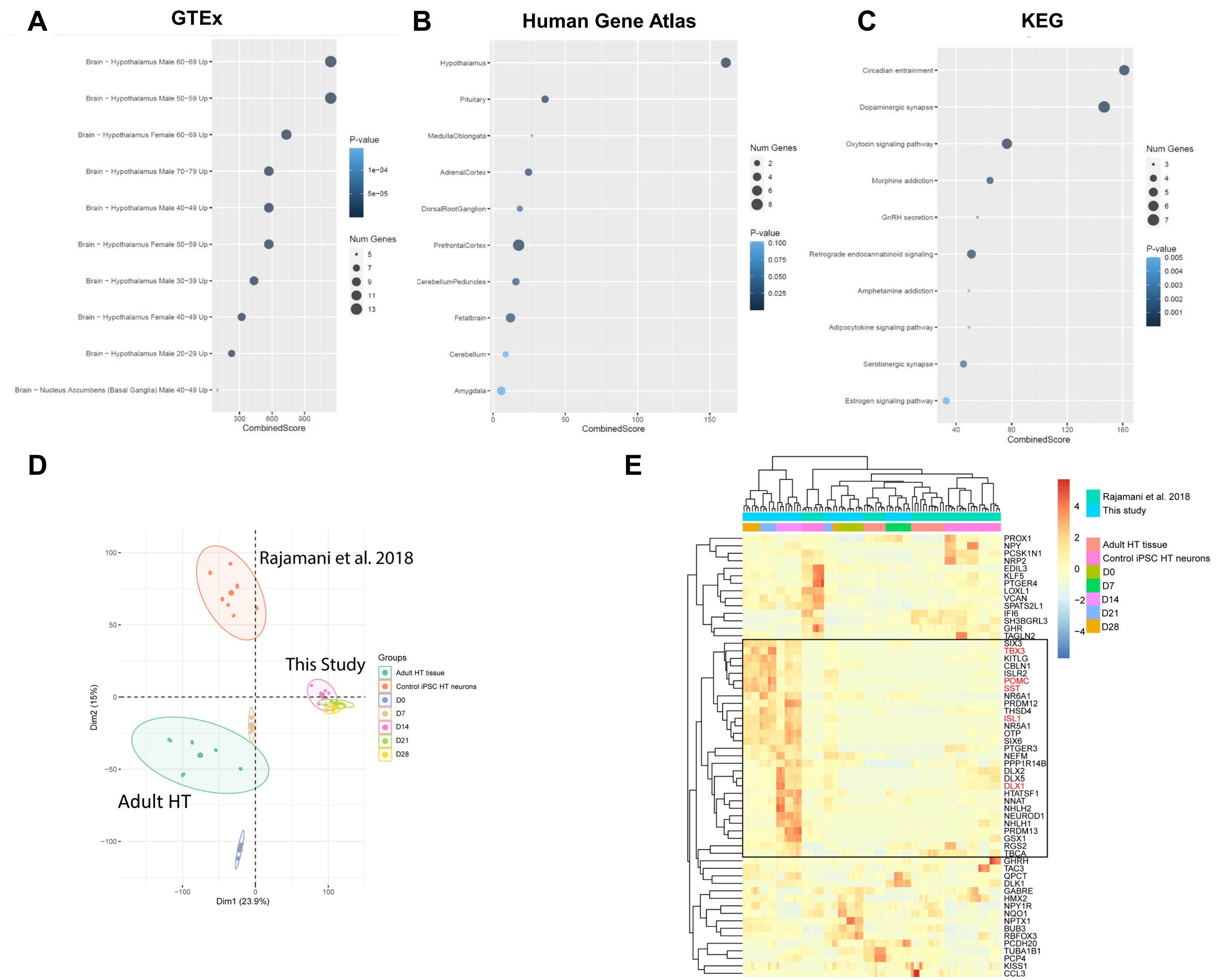

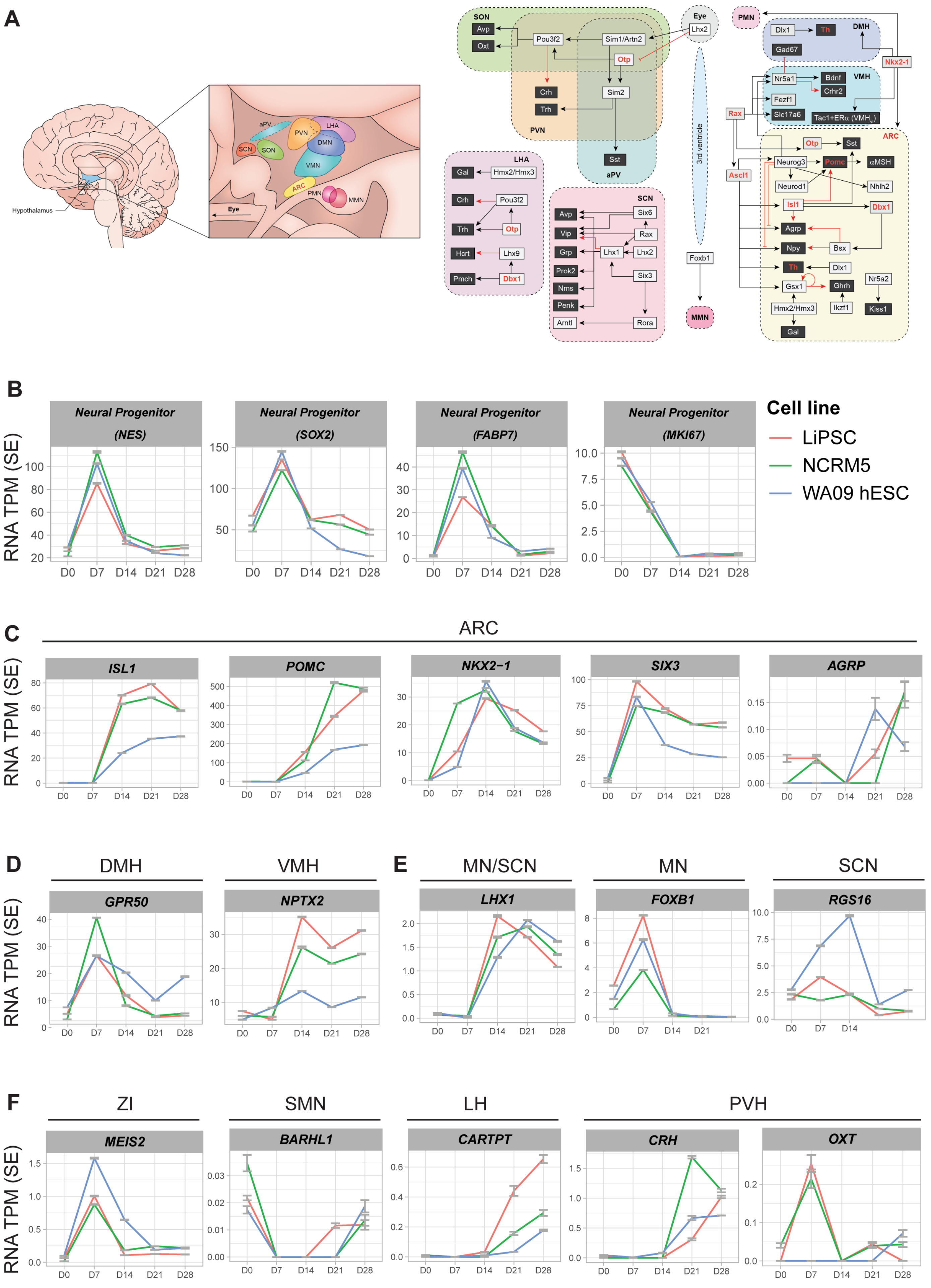

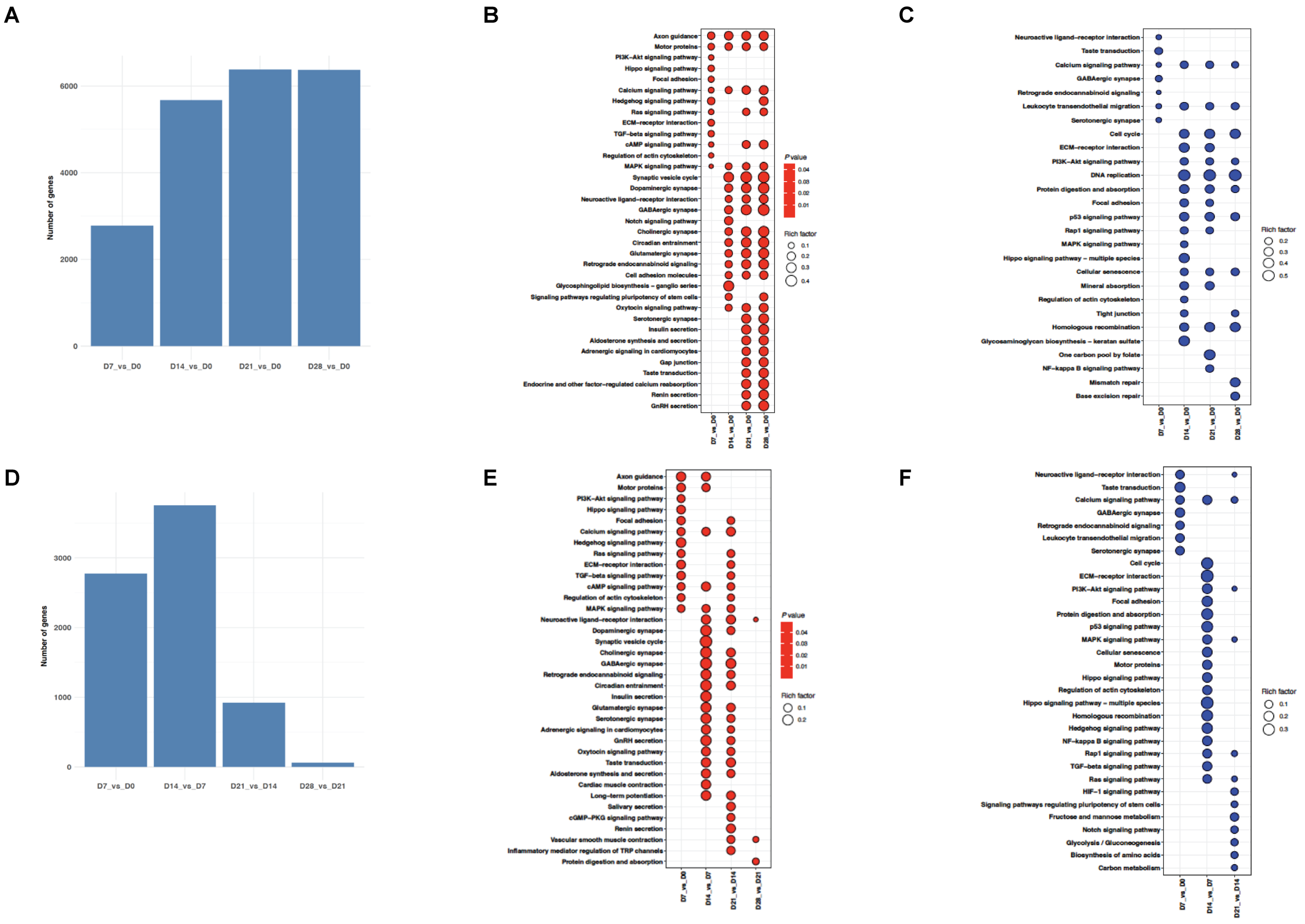
